# Evolutionary repurposing of trypanosomal Pam18 and Pam16 reveals a new regulatory circuit for mitochondrial genome replication

**DOI:** 10.1101/2023.12.05.570232

**Authors:** Corinne von Känel, Silke Oeljeklaus, Salvatore Calderaro, Ignacio M. Durante, Vendula Rašková, Bettina Warscheid, André Schneider

## Abstract

Protein import and genome replication are essential processes for mitochondrial biogenesis and propagation. The J-domain proteins Pam16 and Pam18 regulate the presequence translocase of the mitochondrial inner membrane. In the protozoan *Trypanosoma brucei*, their counterparts are TbPam16 and TbPam18, which are essential for the procyclic form of the parasite, though not involved in mitochondrial protein import. Here, we show that during evolution, the two proteins have been repurposed to regulate the replication of maxicircles within the intricate kDNA network, the most complex mitochondrial genome known. TbPam18 and TbPam16 have inactive J-domains suggesting a function independent of heat shock proteins. However, their single transmembrane domain is essential for function. Pulldown of TbPam16 identifies a putative client protein, termed MaRF11, the depletion of which causes the selective loss of maxicircles, akin to the effects observed for TbPam18 and TbPam16. Moreover depletion of the mitochondrial proteasome results in increased levels of MaRF11. Thus, we propose a model for a membrane-bound regulatory circuit that controls maxicircle replication in response to an unknown nuclear signal. This model posits that MaRF11 directly mediates maxicircle replication, that its level is controlled by proteasomal digestion, and that it is protected from degradation by binding to the TbPam18/TbPam16 dimer.

## Introduction

The parasitic protist *Trypanosoma brucei* has a unique mitochondrial biology. As in other eukaryotes, more than 95% of its mitochondrial proteins are encoded in the nucleus, synthesized in the cytosol and imported into and across the mitochondrial membranes^1^. However, the trypanosomal mitochondrial protein import machineries show significant differences to the prototypical systems of yeast and mammals^2–6^. The largest differences are found in the translocase of the inner mitochondrial membrane, the TIM complex. Essentially all eukaryotes have two TIM complexes, termed TIM22 and TIM23^7,8^. The TIM22 complex mediates insertion of proteins into the inner mitochondrial membrane (IM) that have multi-spanning membrane domains, such as mitochondrial carrier proteins (MCPs)^9,10^. The TIM23 complex imports presequence-containing proteins across or into the IM^11^. To import its substrates into the mitochondrial matrix, TIM23 associates with the matrix-exposed presequence translocase-associated motor (PAM). The PAM consists of five essential and highly conserved subunits^7^: the mitochondrial heat shock protein 70 (mHsp70)^12,13^, its J-domain-containing co-factors Pam18^14–16^ and Pam16^17^, Tim44^18^ and the nucleotide exchange factor Mge1^19–21^.

In contrast, *T. brucei* has a single TIM complex only, which with minor compositional variations, imports both types of substrates^22^. Interestingly, the only trypanosomal TIM component sharing homology to a subunit of TIM complexes in yeast or mammals is TbTim17. TbTim17 is an orthologue of the Tim22 subunit of the TIM22 complex^7,8,23,24^.

To import presequence-containing proteins, the trypanosomal TIM complex associates with a PAM module^42^ containing the trypanosomal mHsp70 orthologue (TbmHsp70), which is essential for the import of presequence-containing proteins^25,26^. *T. brucei* contains *bona fide* orthologues of Pam18 and Pam16, termed TbPam18 and TbPam16. However, while they are essential for normal growth of procyclic form (PCF) trypanosomes, they are not involved in mitochondrial protein import^26^. Instead, the function of Pam18, and likely Pam16, in the trypanosomal PAM is carried out by the non-orthologous, essential, J-domain-containing integral IM protein TbPam27^26^.

Based on these observations, an evolutionary scenario was proposed that aims to explain the transition from two ancestral TIM complexes, found in most eukaryotes, to the single TIM complex of trypanosomes^26^. It posits that in the ancestor of kinetoplastids, TbPam27 fortuitously interacted with TbTim17. This allowed mHsp70 to bind to the resulting TbTim17/TbPam27 complex forming a rudimentary PAM. As a consequence, the TbTim17-containing, TIM22-like TIM complex acquired the capability to import both, presequence-containing proteins and MCPs. Thus, the previously essential TIM23 complex became redundant and its subunits were lost. However, the proposed scenario cannot explain why TbPam18 and TbPam16 were retained during evolution and why they are essential for the growth of PCF *T. brucei*^26^.

The single mitochondrion of trypanosomes contains a single unit genome, termed kinetoplast DNA (kDNA), which is the most complex mitochondrial genome known in nature. The kDNA consists of two types of DNA rings: maxicircles (ca. 25 copies, 23 kb each) and heterogenous minicircles (ca. 5000 copies, 1 kb each) that are arranged in a large intercatenated network^27,28^. Maxicircles encode 16 subunits of the oxidative phosphorylation (OXPHOS) complexes, two mitoribosomal proteins (MRPs) and two rRNAs^27–29^. Twelve of their transcripts require RNA editing to become functional mRNAs.

This process is mediated by small guide RNAs (gRNAs), which are the only genes encoded on the minicircles^30–33^. The kDNA network consists to 90% of minicircles, which are highly topologically interlocked^34^. Maxicircles are also interlocked with each other^35^, and in addition, interwoven into the minicircle network^36^. The resulting kDNA disk in the mitochondrial matrix is physically connected to the flagellum’s basal body in the cytosol via the tripartite attachment complex (TAC)^37,38^.

Minicircle replication begins with their release into the kinetoflagellar zone (KFZ), located between the kDNA disk and the IM^39^. It occurs unidirectionally via theta structures^40^.

Replicated minicircles migrate to the antipodal sites, which are protein complexes at opposing sites of the kDNA disk, where gaps are repaired and minicircles are reattached to the periphery of the network^41,42^. Maxicircles replicate like minicircles, but always remain interlocked with the kDNA. However, the details of the process and the factors required for it are not well understood^28,43^. Minicircle release and reattachment causes concentration of the catenated maxicircles in the center of the disk^35^. The concomitant replication and segregation of the kDNA network, mediated by the TAC and the basal bodies, results in the formation of a maxicircle-containing filament between the two minicircles networks, termed Nabelschnur. Completion of kDNA segregation requires cleavage of this Nabelschnur, to unlink the daughter kDNAs^44,45^.

Altogether, replication of the single kDNA network involves up to 150 different proteins and is tightly coordinated with the nuclear cell cycle^27,28^. However, the mechanism of this coordination is presently unknown. What has been shown is that the mitochondrial proteasome TbHslVU, composed of the two subunits TbHslV and TbHslU, acts a negative regulator of minicircle and maxicircle copy numbers and its depletion thus causes accumulation of giant kDNAs^46,47^. Intriguingly, up to date, only a single TbHslVU substrate has been identified, the maxicircle replication factor TbPIF2, whose levels are increased in TbHslVU depleted cells^47^.

Here, we show that TbPam18 and TbPam16, while not involved in mitochondrial protein import, are required for the replication of the maxicircle component of the kDNA. Strikingly, this function is mediated by a soluble TbPam16-interacting protein whose levels are controlled by TbHslVU.

## Results

### Ablation of TbPam18 or TbPam16 mainly affects MRPs and OXPHOS components

TbPam18 and TbPam16 are not required for protein import, but the fact that they are essential integral IM proteins indicates that they have another function linked to mitochondria^26^. To identify what this function might be, we quantified global changes in the mitochondrial proteome caused by the ablation of either of the two proteins. Previously established tetracycline-inducible TbPam18 and TbPam16 RNAi cell lines^26^ were analyzed by stable isotope labelling by amino acids in cell culture (SILAC)-based quantitative mass spectrometry (MS). Surprisingly, neither TbPam18 nor TbPam16 were detected in the two SILAC RNAi experiments. However, using a TbPam16 antibody, we found that after only one day of RNAi induction, TbPam16 levels were strongly reduced in both cell lines (*Fig. 1A*).

**Figure 1.**
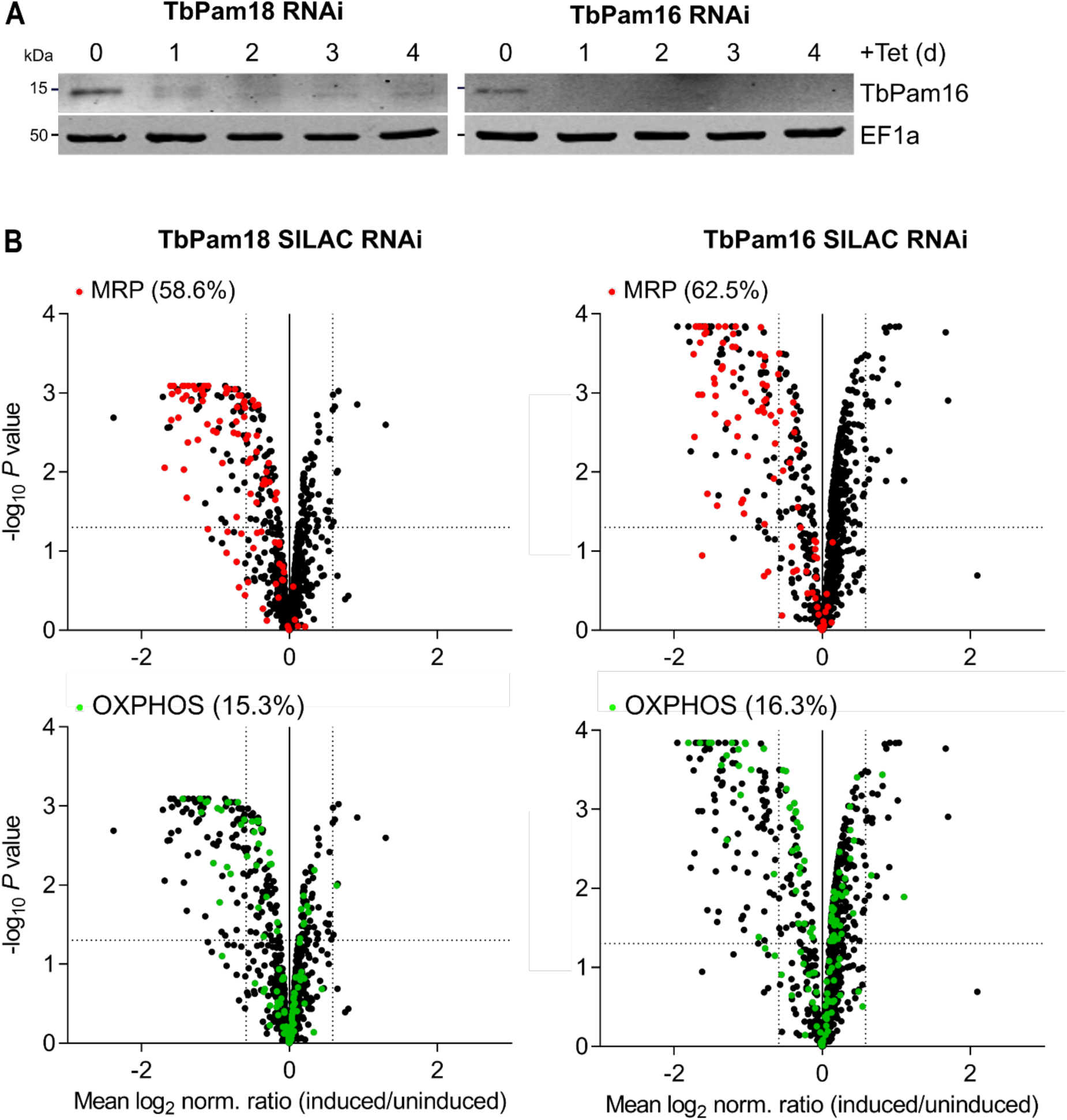
– TbPam18 and TbPam16 RNAi predominantly affects MRPs and OXPHOS components: (A) Immunoblot analysis of steady-state protein levels of TbPam16 in whole-cell extracts of TbPam16 or TbPam18 RNAi cell lines over four days of induction. EF1a serves as loading control. **(B)** Global mitochondrial proteome changes upon ablation of TbPam18 (leÉ panels) or TbPam16 (right panels). Mitochondria-enriched fractions of uninduced and four days induced TbPam18 and TbPam16 RNAi cells were analyzed by SILAC-based quantitative mass spectrometry. Datasets were filtered for mitochondrial proteins and the mean log_2_ of normalized ratios (induced/uninduced) was ploÖed against the corresponding negative log_10_ of the corrected and adjusted *P* value (limma test). Highlighted are mitochondrial ribosomal proteins (MRPs, red) and components of the oxidative phosphorylation pathway (OXPHOS, green). The horizontal doÖed line in each volcano plot marks an adjusted *P* value of 0.05. The vertical doÖed lines indicate a fold-change in protein abundance of ±1.5.

Thus, we conclude that i) the stability of TbPam16 depends on TbPam18, suggesting the two proteins form a heterodimer as in yeast, and ii) that RNAi against TbPam16 is very efficient.

916 and 893 mitochondrial proteins^1,29,48^ were detected in the SILAC RNAi datasets and the levels of 12.4% and 13.2% of them were reduced more than 1.5-fold in the TbPam18 and TbPam16 RNAi cell lines, respectively (*Fig. 1B*). The most affected proteins included mitoribosomal proteins (MRPs)^3^, of which 58.6% and 62.5% were depleted more than 1.5-fold in the two cell lines (*Fig. 1B, top panels*). Furthermore, we found that 15.3% and 16.3%

of all detected components of the OXPHOS pathway^48^ were reduced more than 1.5-fold in the two cell lines (*Fig. 1B, bottom panels*). In both experiments, complex IV was affected the most, followed by complexes I and III, whereas the levels of complex II and V subunits were not or only marginally decreased.

A common feature of the mitoribosome and the OXPHOS complexes I, III and IV is that some of their subunits are encoded on the kDNA^27,29,49^. For mitoribosomes, these are the 12S and 9S rRNAs as well as two MRPs^29^. For complexes I, III and IV the number of maxicircle-encoded subunits is six, three and three, respectively. In contrast, only a single complex V subunit is encoded on the kDNA and all complex II subunits are encoded in the nuclear genome^49^.

Taken together, the SILAC RNAi results presented here indicate that the ablations of TbPam18 and TbPam16 could affect the kDNA.

### Depletion of TbPam18 and TbPam16 leads to the loss of maxicircles

To investigate the fate of the kDNA upon TbPam18 or TbPam16 depletion, we analyzed DAPI-stained RNAi cells by fluorescence microscopy. In line with the SILAC-RNAi analyses (*Fig 1B*), we found that after four days of RNAi induction, many TbPam18 and TbPam16 RNAi cells had smaller kDNAs compared to uninduced cells (*Fig. 2A, upper panels*). A quantification of the experiments (*Fig. 2A, lower panels*) showed in both cell lines a time-dependent decrease of the kDNA size to 90% and about 60% after three (prior to the onset of the growth retardation) to five days of RNAi induction, respectively. Shrinkage of the kDNA disk has been observed previously when proteins involved in kDNA replication were ablated^47,50–54^.

**Figure 2.**
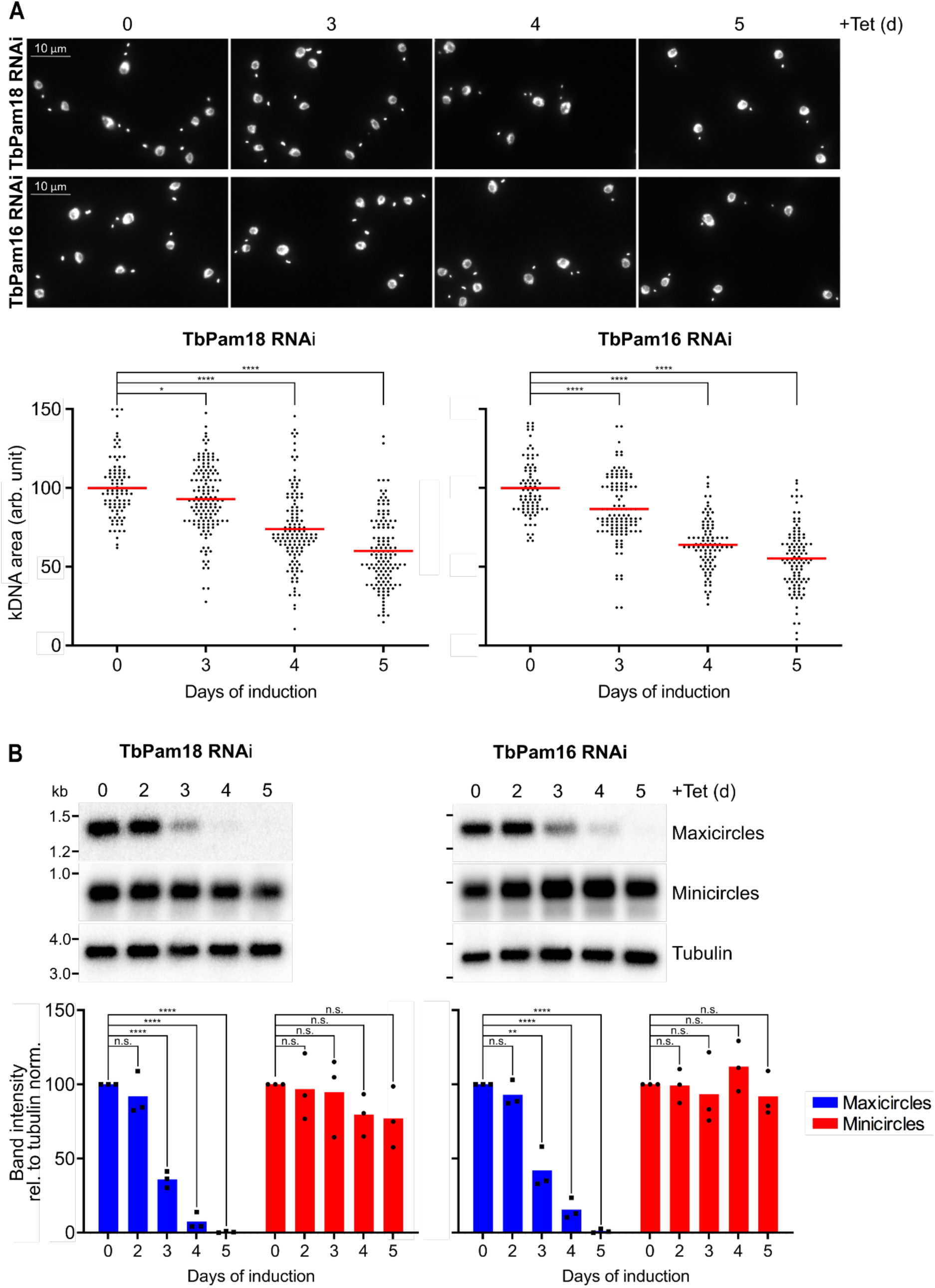
– TbPam18 and TbPam16 abla_on causes the loss of maxicircles: (A) Upper panels: Fluorescence microscopy analysis of DAPI-stained uninduced and three to five days induced TbPam18 and TbPam16 RNAi cells. Lower panels: Quantification of kDNA areas in 86 to 140 DAPI-stained RNAi cells induced for the indicated amount of Çme. The red line indicates the mean of the kDNA areas at each timepoint. The mean of the uninduced cells was set to 100%. *: *P* value<0.05, ****: *P* value<0.0001, as calculated by an unpaired two-tailed t-test. **(B)** Southern blot analysis of steady-state levels of mini– and maxicircles in the TbPam18 and TbPam16 RNAi cell lines. Upper panels: Total DNA from uninduced or three to five days induced cells was isolated and digested with HindIII and XbaI. Probes specifically recognizing mini– or maxicircles were used. A probe detecting a 3.6-kb fragment of the tubulin intergenic region serves as loading control. Lower panels: Densitometric quantification of mini– and maxicircle abundance on Southern blots. The ratio of the mini– or maxicircle abundance and the respective loading control (tubulin) was normalized (norm.) to the ratios of uninduced cells. Blue (maxicircles) and red (minicircles) bars represent the mean of three independent biological replicates. n.s.: not significant, **: *P* value<0.01, ****: *P* value<0.0001, as calculated by an unpaired two-tailed t-test.

To study the effects on the kDNA in more detail, we performed Southern blot analysis (*Fig. 2B*). Total DNA was extracted from uninduced and induced RNAi cells, digested by restriction enzymes and separated on an agarose gel. The resulting blot was hybridized with mini– and maxicircle specific probes. Already after three days of RNAi induction, maxicircle levels are significantly reduced to about 39% in both cell lines. After five days they are almost undetectable. In contrast, the levels of minicircles are not significantly changed over five days of RNAi induction. The same experiment (*Fig. 2B*) was repeated using PCR, to detect the changes in mini– and maxicircle levels and the same results were obtained (*Fig. S1)*. The observation that the depletion of maxicircles is detected prior to the onsets of growth retardations (which occur at day four)^26^ suggests that TbPam18 and TbPam16 are directly involved in maxicircle replication or maintenance.

Since minicircles make up 90% of the kDNA^34^, the massive network shrinkage seen in the DAPI stains of *Fig. 2A* cannot be explained by a selective loss of maxicircles only ^1,8^. One way to explain the constant levels of minicircles during TbPam18 and TbPam16 depletion (*Fig. 2B*) is that they are released from the kDNA disk that progressively gets depleted from maxicircles.

To detect potential changes in free minicircle levels, digitonin-extracted, mitochondria-enriched pellets from uninduced and induced TbPam18 and TbPam16 RNAi cells were solubilized in 1% digitonin. Subsequent centrifugation resulted in pellets containing kDNA networks and supernatants containing free minicircles. PCR analysis of the DNA extracted from these fractions showed that maxicircles were only present in the pellets and that their levels decreased over time of induction as expected (*Fig 3A*). Minicircles behaved very differently. In uninduced cells they were almost exclusively found in pellet fractions and thus in the kDNA networks. However, during the course of the RNAi, the amount of detected minicircles completely shifts to the supernatant (*Fig. 3A*). Thus, ablation of TbPam18 and TbPam16, and the accompanying loss of maxicircles, does not inhibit the release of minicircles from the remaining kDNA network, nor their replication. But it appears to prevent their reattachment to the maxicircle-depleted networks (*Fig. 3B*).

**Figure 3.**
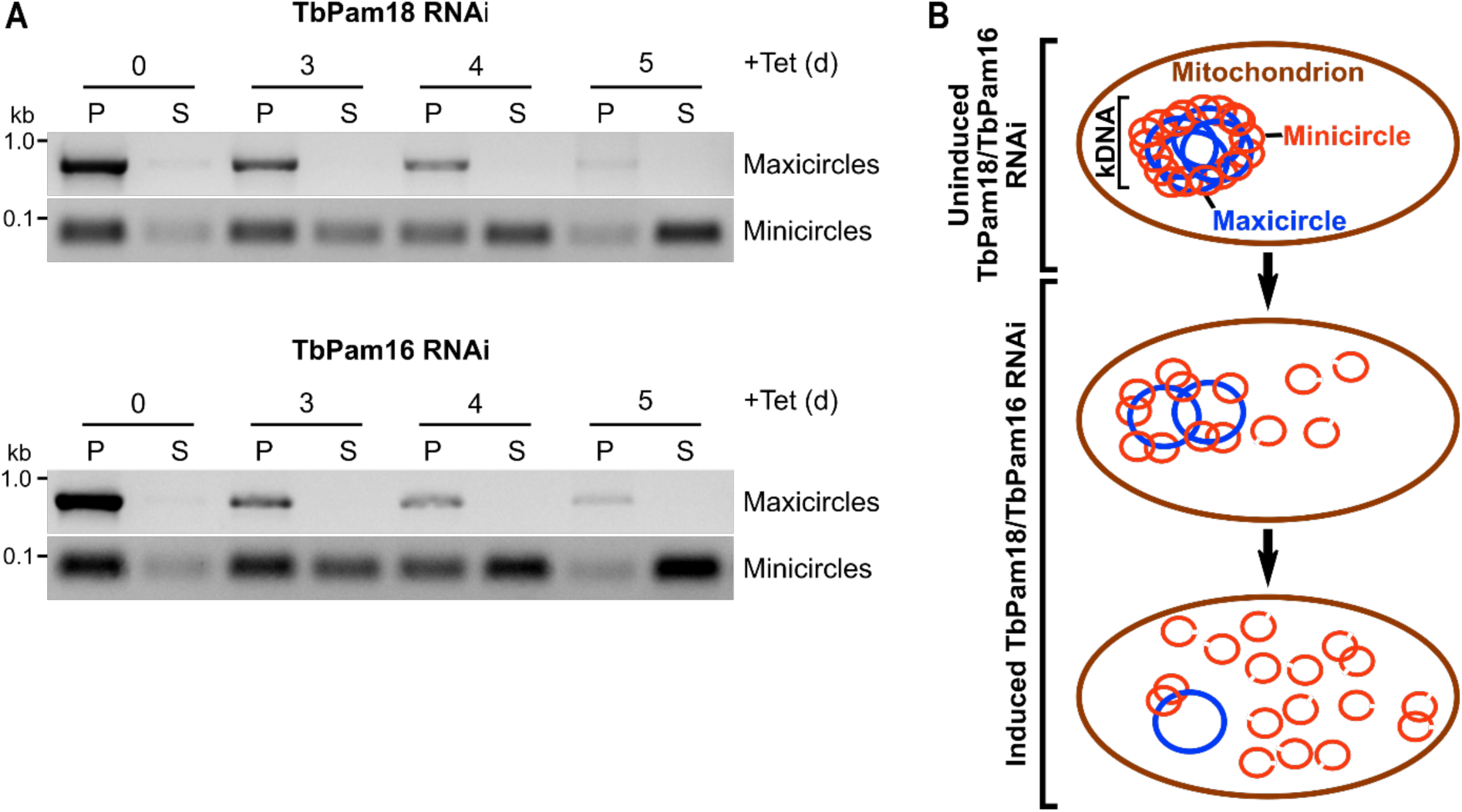
– Abla_on of TbPam18 and TbPam16 causes accumula_on of free minicircles: (A) A quantitative PCR-based method was used to analyze steady state levels of kDNA-bound or free mini– and maxicircles. Digitonin-extracted, mitochondria-enriched pellets from uninduced and three to five days induced TbPam18 and TbPam16 RNAi cells were solubilized in 1% digitonin. Subsequent centrifugation resulted in a pellet fraction (P) containing intact kDNA networks and a soluble fraction (S) containing free minicircles. DNA extracted from both fractions was used as template for PCR reactions amplifying specific mini– or maxicircle regions. PCR products were analyzed on agarose gels. **(B)** Schematic illustration of the putative sequence of effects on mini– and maxicircles upon RNAi-induced knockdown of TbPam18 and TbPam16. The ablation of TbPam18 and TbPam16 and the concomitant loss of maxicircles does not seem to inhibit the release of minicircles from the kDNA, nor their replication. However, it appears to prevent the reaÖachment of free minicircles to the kDNA network. Consequently, free minicircles accumulate in the mitochondrial matrix.

### TbPam18 and TbPam16 have procyclic form-specific functions

*T. brucei* has a complex life cycle alternating between an insect vector, the Tsetse fly, and a mammalian host. One of the replicative stages in the insect vector is the PCF, which contains an extensively reticulated mitochondrion that is capable of OXPHOS. The replicative stage in the mammalian host is the bloodstream form (BSF), which has a less reticulated mitochondrion that cannot perform OXPHOS^27,48^. The BSF produces its energy exclusively by glycolysis. But because the mitochondrial membrane potential in BSFs is maintained by the F1Fo ATP synthase working in reverse, a subunit of which is encoded on the kDNA, an intact kDNA network is essential not only in the PCF but also for the BSF^55,56^.

It was therefore surprising that RNAi-mediated ablation of TbPam18 in the New York single marker (NYsm) BSF strain^57^ did not affect growth (*Fig. 4A*). However, the interpretation of this result is complicated, because even efficient RNAi never eliminates all mRNAs. It could be that the small amount of TbPam18 still present in these cells is sufficient for growth. To not run into the same issue with TbPam16, we established a TbPam16 NYsm double knockout (dKO) cell line, whose growth was indistinguishable from its parent cell line (*Fig. 4B*). This demonstrates that the function of TbPam16 is indeed specific for PCF trypanosomes and that its role in maxicircle replication is redundant in BSF cells or taken over by another protein. For unknown reason we were not able to produce a dKO cell line for TbPam18, not even a conditional one. However, based on the results of the TbPam18 RNAi (*Fig. 4A*) and the observation that TbPam16 and TbPam18, as Pam18 and Pam16 in yeast, likely form a heterodimer (*Fig. 1A*), we conclude that the function of TbPam18 is also redundant in the BSF.

**Figure 4.**
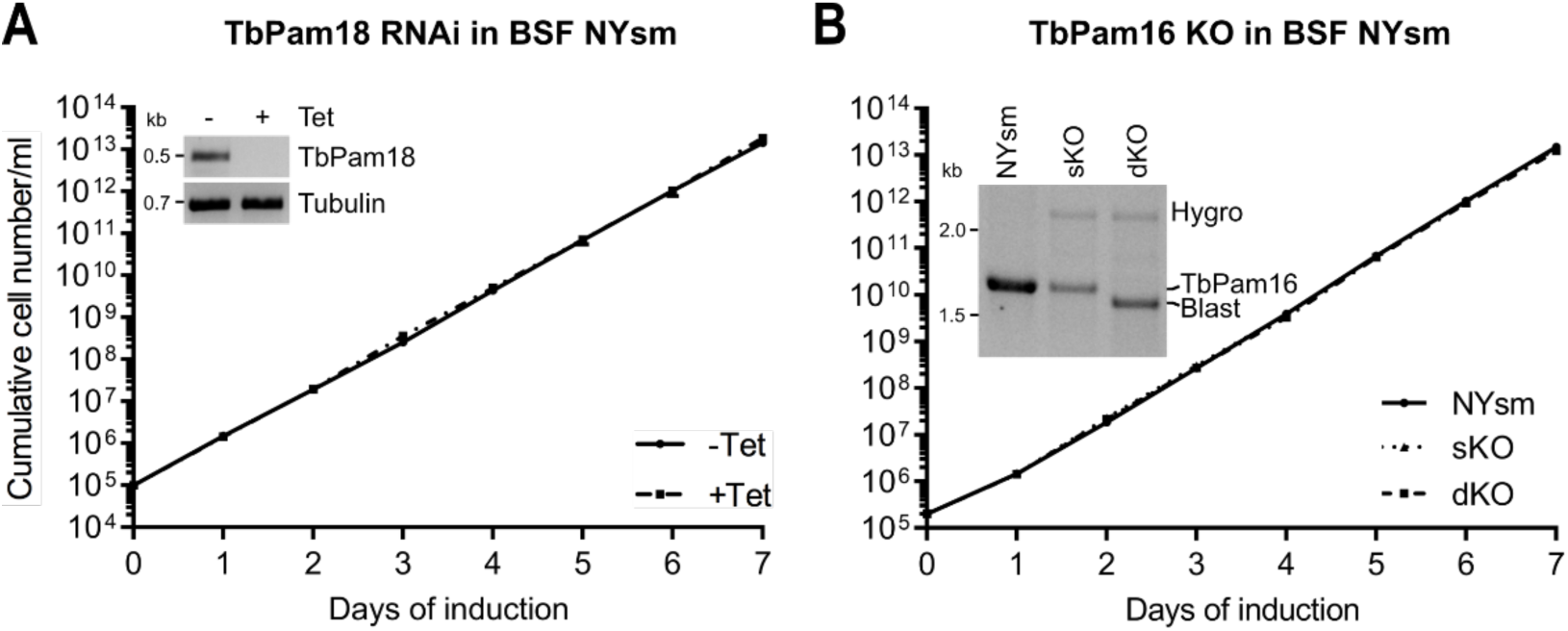
– TbPam18 and TbPam16 are not essen_al in BSF trypanosomes: (A) Growth curve of uninduced (-Tet) and induced (+Tet) bloodstream form (BSF) New York single marker (NYsm) RNAi cell line ablating TbPam18. Error bars correspond to the standard deviation (n=3, error bars are too small to be visible). Inset: RT-PCR product of the wt TbPam18 mRNA in uninduced (-) or two days induced (+) cells. Tubulin mRNA serves as loading control. **(B)** Growth curve of NYsm, TbPam16 single knockout (sKO) and double knockout (dKO) BSF cell lines. Inset: Verification of sKO and dKO by PCR using one primer pair to amplify the TbPam16 ORF (∼1.7 kilobases (kb)), the hygromycin (hygro, ∼2.3 kb) or blasticidin (blast, ∼1.6 kb) resistance casseÖes at the same Çme. Hygro was used to replace the first allele and blast was used to replace the second allele.

### Integral membrane localization of TbPam18 and TbPam16 is functionally relevant

TbPam18 and TbPam16 are integral IM proteins^26^, which raises the question whether this feature is essential for their function. To find out, we expressed RNAi-resistant (RNAi-res.) TbPam18 and TbPam16 variants lacking their predicted transmembrane domains^58^ (TMDs) as well as their short IMS exposed N-termini (4 aa for TbPam18 and 33 aa for TbPam16) (*Fig. 5A*)^55^. Note that only the TbPam16 variants could be C-terminally HA-tagged, because in the case of TbPam18, both N– and C-terminal tags abolished the function of the protein (*Fig. 5A, Fig. S2A*). Expression of full-length TbPam18 or TbPam16-HA in RNAi cells ablated for the corresponding endogenous proteins, restored normal growth as expected (*Fig. 5BC, left panels*). However, the same was not the case for the ΔN-TbPam18 and ΔN-TbPam16-HA variants that lack the predicted N-terminal TMDs (*Fig. 5BC, right panels*). Both truncated variants were N-terminally fused to the mitochondrial targeting sequence (MTS) of trypanosomal TbmHsp60, to ensure their import into mitochondria. Since the TbPam16-HA and ΔN-TbPam16-HA variants were tagged, their import could be verified biochemically using digitonin extractions (*Fig. S2B*). Moreover, alkaline carbonate extractions showed that as expected for an integral membrane protein, full-length TbPam16-HA is exclusively recovered in the pellet. In contrast, the major fraction of the truncated ΔN-TbPam16-HA is recovered in the supernatant indicating it is soluble.

**Figure 5.**
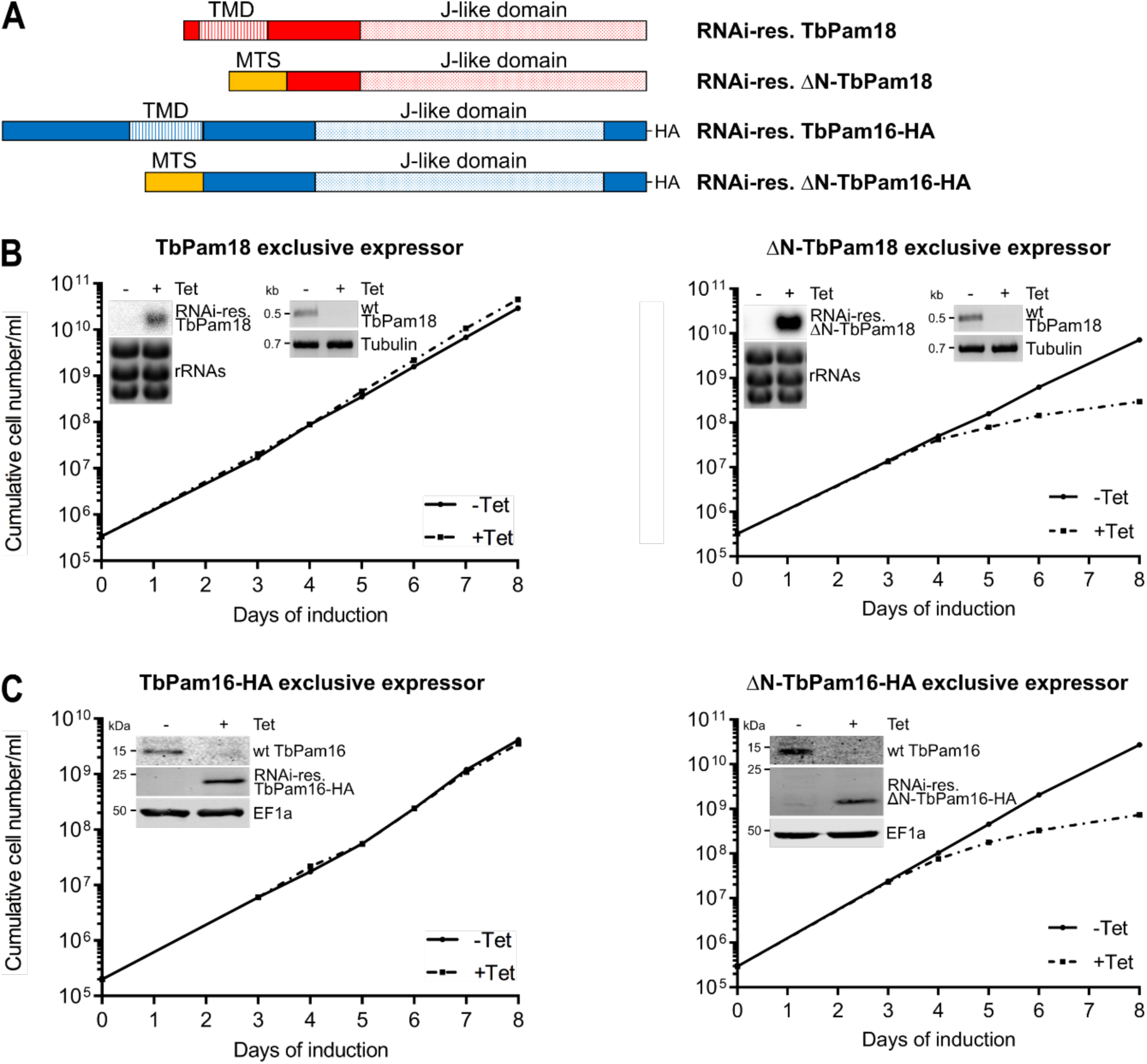
– Integral membrane localiza_on is crucial for TbPam18’s and TbPam16’s func_ons: (A) Schematic representation of RNAi-resistant (RNAi-res.) full-length (TbPam18 and TbPam16-HA) and N–terminally truncated variants of TbPam18 (ΔN-TbPam18) and TbPam16 (ΔN-TbPam16-HA). TbPam18 constructs are untagged, while TbPam16 constructs carry a C-terminal HA-tag. Predicted transmembrane domains (TMD) and J-like domains are indicated. To ensure mitochondrial localization, the N-terminally truncated variants were expressed carrying the mitochondrial targeting sequence (MTS) of trypanosomal mitochondrial heat shock protein 60. **(B)** Growth curves of uninduced (-Tet) and induced (+Tet) cell lines ectopically expressing RNAi-res., full-length TbPam18 (leÉ) or ΔN– TbPam18 (right) in the background of RNAi targeting the wildtype (wt) TbPam18 (TbPam18 and ΔN-TbPam18 exclusive expressors). Insets on the leÉ: Northern blots of total RNA isolated from uninduced (-) and two days induced (+) cells probed for the mRNAs of RNAi-res. TbPam18 or ΔN-TbPam18 to confirm effcient inducible ectopic expression. Ethidium bromide-stained rRNAs serve as loading control. Insets on the right: RT-PCR products of the wt TbPam18 mRNA in uninduced (-) or two days induced (+) cells. Tubulin mRNA serves as loading control. **(C)** Growth curve of uninduced (-Tet) and induced (+Tet) cells ectopically expressing RNAi-res. TbPam16-HA (leÉ) or ΔN-TbPam16-HA (right) in the background of RNAi targeting the wt TbPam16 (TbPam16-HA and ΔN-TbPam16-HA exclusive expressors). Insets: Immunoblot analysis of whole-cell extracts of uninduced (-) and two days induced (+) cells probed for wt TbPam16, RNAi-res. TbPam16-HA or ΔN-TbPam16-HA and EF1a as loading control.

The results in *Fig. 5* suggest that the integration of TbPam18 and TbPam16 into the IM is essential for their function and thus link maxicircle replication to the IM.

### The J-domain of yeast Pam18 cannot complement the loss of the TbPam18 J-like domain

Classic Pam18 homologues involved in mitochondrial protein import contain the highly conserved tripeptide HPD in their J-domain (*Fig. S3A*)^57^, which is essential for the stimulation of the ATPase activity of their Hsp70 partners^59,60^. In contrast, TbPam18 has a degenerate J-domain containing the tripeptide HSD making it a J-like protein^60^, a feature that is highly conserved within kinetoplastids (*Fig. S3B*). Thus, we wondered i) whether the HSD motif of TbPam18 is important for its function and ii) if the J-domain of yeast Pam18 (ScPam18) can take over the function of the J-like domain of TbPam18.

To that end, we generated a cell line allowing the exclusive expression of a TbPam18 variant in which the tripeptide HSD was mutated to HPD (TbPam18-S98P, *Fig 6A*).

**Figure 6.**
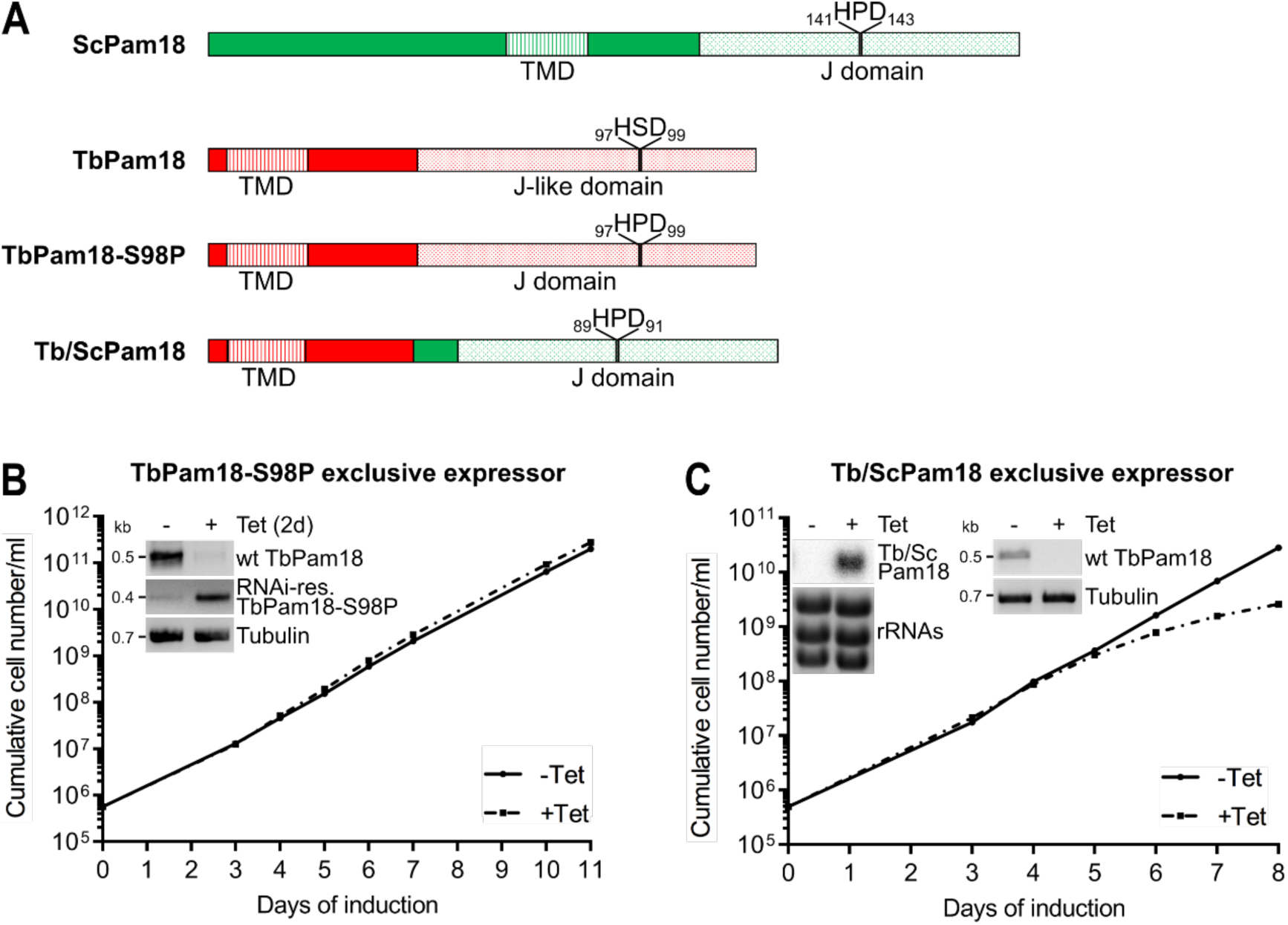
– The J-domain of ScPam18 cannot complement the loss of the J-like domain of TbPam18: (A) Schematic representations of yeast (Sc) Pam18, TbPam18 and a mutated TbPam18 version, in which the J-like domain was altered to a J-domain by changing the serine residue at position 98 to a proline residue (TbPam18-S98P). Finally, a Tb/Sc fusion Pam18 (Tb/ScPam18), in which the J-like domain of TbPam18 was replaced by the J-domain of ScPam18 is shown. **(B)** Growth curve of uninduced (-Tet) and induced (+Tet) cells ectopically expressing RNAi-res. TbPam18-S98P in the background of RNAi targeting the endogenous wt TbPam18 (TbPam18-S98P exclusive expressor). Inset: RT-PCR products of wt TbPam18 and TbPam18-S98P mRNAs in uninduced (-) or two days induced (+) cells. Tubulin mRNA serves as loading control. **(C)** Growth curve of uninduced (-Tet) and induced (+Tet) cells ectopically expressing RNAi-resistant (RNAi-res.) Tb/ScPam18 in the background of RNAi targeting the endogenous wildtype (wt) TbPam18 (Tb/ScPam18 exclusive expressor). Inset on the leÉ: Northern blot of total RNA extracted from uninduced (-) and two days induced (+) cells, probed for Tb/ScPam18, to confirm inducible ectopic expression. Inset on the right: RT-PCR product of the wt TbPam18 mRNA in uninduced (-) or two days induced (+) cells. Tubulin mRNA serves as loading control.

Intriguingly, this variant can fully complement the growth retardation caused by the RNAi-mediated depletion of TbPam18 (*Fig. 6B*). Thus, TbPam18 can function with both, J or J-like domains.

To find out whether the intact J-domain of ScPam18 can replace the J-like domain of TbPam18, we used a chimeric protein consisting of the TMD of TbPam18 and the J-domain of ScPam18 (Tb/ScPam18) (*Fig. 6A*). Expression of Tb/ScPam18 in TbPam18 RNAi background delayed the onset of the growth phenotype by one day but could not rescue the growth retardation at later time points (*Fig. 6C*).

Because tagged versions of TbPam18 are not functional (*Fig. S2A*), the variants tested above were untagged. Nevertheless, we analyzed the localization of N-terminally myc-tagged TbPam18 and Tb/ScPam18, which showed that about half of each variant is recovered in the mitochondria-enriched fraction of a digitonin extraction *(Fig. S3C)*.

Moreover, in an alkaline carbonate extraction, both TbPam18 versions present in the mitochondria-enriched fraction are exclusively recovered in the pellet indicating they are integrated into the IM. This strongly suggest that also untagged TbPam18 and Tb/ScPam18 are correctly localized.

In summary, these results show that some feature of the ScPam18 J-domain, other than the HSD or HPD tripeptide, is incompatible with the function of TbPam18.

### TbPam16 interacts with TbPam18 and two additional essential proteins

We have previously used TbPam18-HA and TbPam16-HA for SILAC co-immunoprecipitations experiments (CoIP)^26^. However, tagged TbPam18 is not functional (*Fig. S2A*). Moreover, in the TbPam16-HA CoIP experiment, the only interactor identified was TbPam18^26^. Thus, in hindsight, these experiments are difficult to interpret. Therefore, we repeated the TbPam16 SILAC CoIP with two modifications. First, we used the newly generated cell line allowing exclusive expression of functional TbPam16-HA (*Fig. 5C*), and second, we analyzed both premix as well as postmix samples. Premix conditions means that differentially labelled uninduced and induced cells are mixed prior to the CoIP, which preferentially detects stable interaction partners. In contrast, in the postmix sample, the eluates from separately generated, differentially labeled CoIPs are mixed, allowing the detection of both stable and more transient interaction partners^61^.

In the premix experiment, TbPam16 has been enriched 5.3-fold and in the postmix experiment 8.1-fold demonstrating that both CoIPs were successful *(Fig. 7A)*. Importantly, apart from the bait TbPam16, the most enriched protein in both experiments was TbPam18 confirming the interaction between the two proteins.

**Figure 7.**
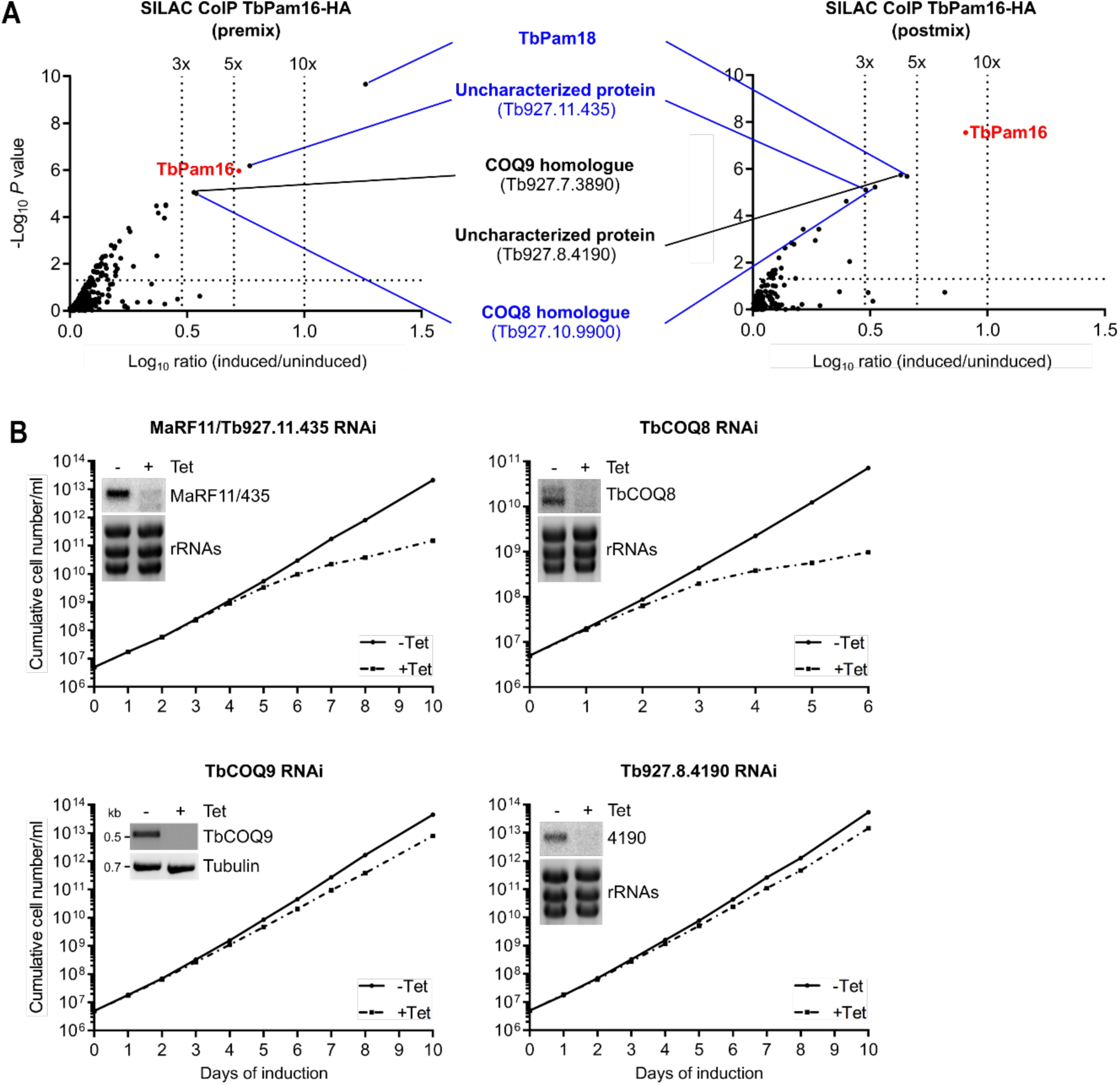
– TbPam16 interacts with TbPam18 and two other essen_al proteins: (A) Volcano plots depicting proteins detected in SILAC-based quantitative mass spectrometry analysis of TbPam16-HA CoIPs. In the experiment on the leÉ, differentially labelled uninduced and induced cells were mixed and the resulting mixture was subjected to CoIP (premix). In the experiment on the right, CoIPs with uninduced and induced cells were done separately and the resulting eluates were mixed aÉerwards (postmix). The vertical doÖed lines in the volcano blots indicate the specified enrichment factors. The horizontal doÖed line indicates a rank-sum test significance level of 0.05. The bait TbPam16 is highlighted in red. Proteins that were significantly detected and enriched more than three-fold in the pre-as well as the postmix experiments are labelled in blue. Proteins enriched more than three-fold in either the pre-or the postmix experiment are labelled in black. **(B)** Growth curves of uninduced (-Tet) and induced (+Tet) MaRF11/Tb927.11.435, TbCOQ8, TbCOQ9 and Tb927.8.4190 RNAi cells. Insets in the MaRF11/Tb927.11.435, TbCOQ8 and Tb927.8.4190 RNAi growth curves: Northern blots of total RNA extracted from uninduced (-) and two days induced cells (+), probed for the respective mRNAs. Ethidium bromide-stained ribosomal RNAs (rRNAs) serve as loading controls. Inset in the TbCOQ9 RNAi growth curve: RT-PCR product of the TbCOQ9 mRNA in uninduced (-) or two days induced (+) cells. Tubulin mRNA serves as loading control.

Interestingly, in addition to TbPam18, two other proteins were enriched more than threefold in both experiments (*Fig. 7A*). The first one is a trypanosome-specific, uncharacterized protein (Tb927.11.435) with a molecular weight of only 10.6 kDa and a remarkably high isoelectric point (pI=10.6). This is interesting because a high pI is a characteristic often found in DNA binding proteins. An AlphaFold structure prediction of Tb927.11.435 reveals that it consists of four consecutive in part amphiphilic α-helices (*Fig S4*)^62^. The second protein, Tb927.10.9900 (*Fig. 7A*), is homologous to yeast and human COQ8 and thus was termed TbCOQ8. COQ8 is a subunit of the coenzyme Q biosynthetic complex (complex Q), located on the matrix face of the IM^63^. Intriguingly, in the TbPam16-HA premix SILAC CoIP, Tb927.7.3890, a protein orthologous to complex Q subunit 9 (COQ9) was detected. The presence of TbCOQ8 and TbCOQ9 amongst the most enriched proteins in the TbPam16-HA SILAC CoIPs is striking. However, only little is known about ubiquinone biosynthesis in *T. brucei* and neither of the two subunits have been previously analyzed. In the postmix experiment, the fifth protein enriched more than threefold is Tb927.8.4190, another trypanosome-specific, uncharacterized protein.

To further study the four newly identified TbPam16 interaction partners, we generated RNAi cell lines. Growth curves revealed that knockdowns of Tb927.11.435 and TbCOQ8 cause a retardation in growth starting after five and two days of induction, respectively (*Fig. 7B*). In contrast, RNAi against TbCOQ9 and Tb927.8.4190 only marginally affected growth. Hence, we focused on Tb927.11.435 and TbCOQ8 in further experiments.

### Tb927.11.435 ablation phenocopies the depletions of TbPam18 and TbPam16

Next, we investigated the fate of the kDNA upon Tb927.11.435 or TbCOQ8 depletion using the same methods that were applied for TbPam16 and TbPam18 (*Fig. 2A, 3 and Fig. S3A*).

Quantification of DAPI-stained RNAi cells using fluorescence microscopy showed that in the Tb927.11.435 RNAi cell line the kDNA size was significantly reduced to 80% and 36% after three (prior to the onset of the growth retardation at day four) to six days of RNAi induction, respectively (*Fig. 8A*). On the contrary, ablation of TbCOQ8 was not found to significantly change the size of the kDNA over four days of RNAi induction. The kDNA network in the Tb927.11.435 RNAi cell line was further analyzed by quantitative PCR, which showed a decrease in the amount of maxicircles to 31% and less than 10% after three to four days of RNAi induction, respectively. In contrast, there was no significant change in the amount of minicircles for at least six days (*Fig. 8B*). However, while the total amount of minicircles remained constant, they were progressively and essentially completely released from the kDNA network within five days of induction (*Fig. 8C*). Thus, depletion of Tb927.11.435 exactly phenocopied the results observed in induced TbPam16 and TbPam18 PCF RNAi cells, which is why we named it maxicircle replication factor of 11 kDa (MaRF11) (*Fig. 2, 3 and Fig. S3.*1). Furthermore, this also holds true for RNAi-induced knockdown of MaRF11 in BSF NYsm cells, which does not cause a change in the growth rate (*Fig. 4, Fig. 8D*).

**Figure 8.**
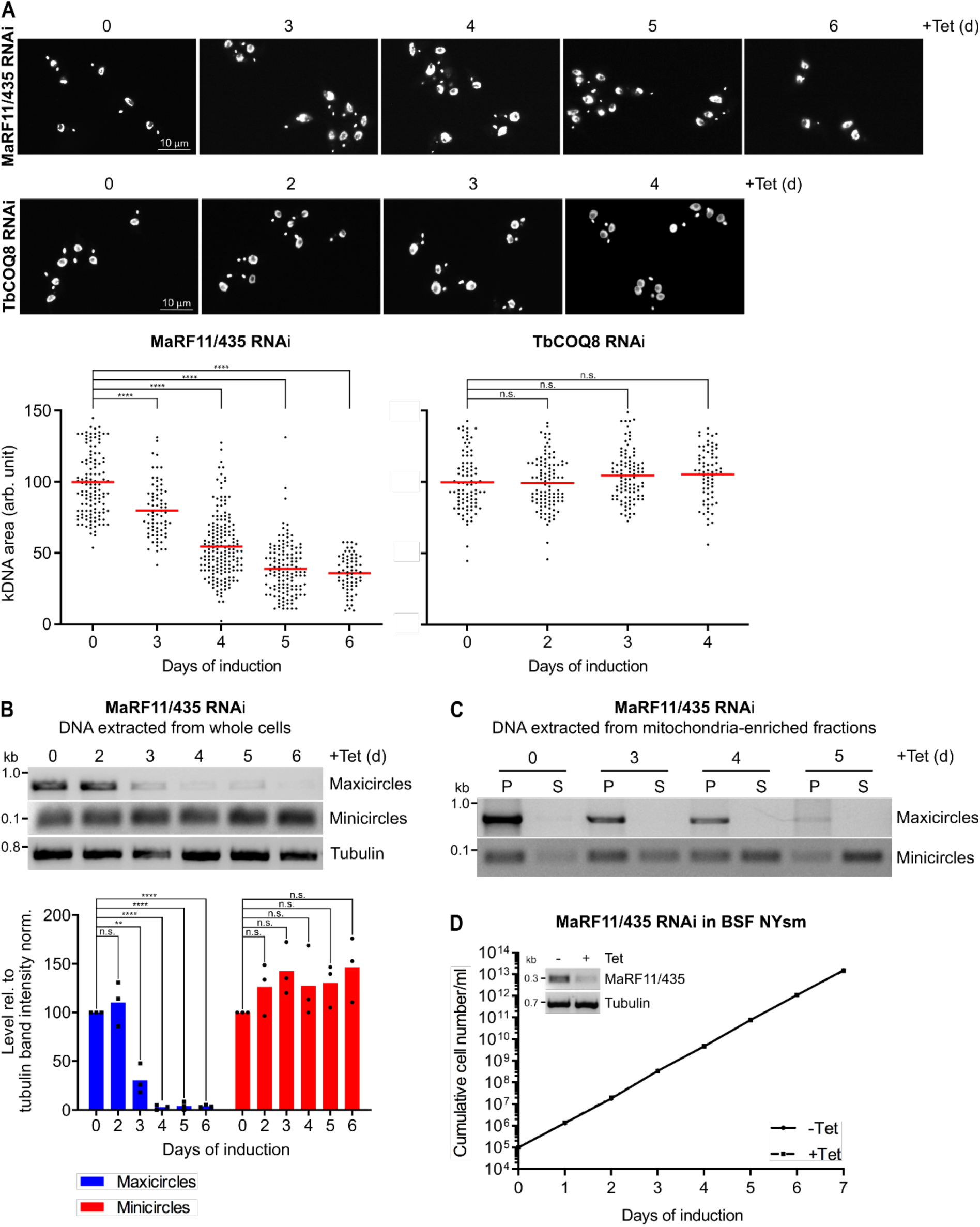
– Abla_on of MaRF11/Tb927.11.435 phenocopies the deple_ons of TbPam18 and TbPam16 in PCF and BSF: (A) Upper panels: Fluorescence microscopy analysis of DAPI-stained uninduced and three to six days induced MaRF11/Tb927.11.435 (435) RNAi cells as well as uninduced and two to four days induced TbCOQ8 RNAi cells. Lower panels: Quantification of kDNA areas in 65 to 175 DAPI-stained RNAi cells. Red line indicates the mean of the kDNA size at each timepoint. The mean of the uninduced cells was set to 100%. n.s.= not significant, ****: *P* value<0.0001, as calculated by an unpaired two-tailed t-test. **(B)** Upper panel: A quantitative PCR-based (qPCR) method was used to detect changes in steady-state levels of total mini– and maxicircles. Total DNA was extracted from uninduced and two to six days induced MaRF11/435 RNAi cells. This DNA was used as the template in PCRs amplifying specific mini– and maxicircle regions or the intergenic region of tubulin. PCR products were analyzed on agarose gels. Lower panels: Densitometric quantification of mini– and maxicircle abundance as detected by qPCR. The ratio of the mini– or maxicircle band intensity and the respective loading control (tubulin band intensity) was normalized (norm.) to the ratios of uninduced cells. Blue (maxicircles) and red (minicircles) bars represent the mean of three independent biological replicates. n.s.: not significant, **: *P* value<0.05, ****: *P* value<0.0001, as calculated by an unpaired two-tailed t-test. **(C)** A quantitative PCR-based method used to analyze steady-state levels of kDNA-bound or free mini– and maxicircles. A digitonin-extracted, mitochondria-enriched pellet from uninduced and three to five days induced MaRF11/435 RNAi cells was solubilized in 1% digitonin. A subsequent centrifugation step resulted in a pellet fraction (P) containing intact kDNA networks and a soluble fraction (S) containing free minicircles. DNA extracted from both fractions was used as template for PCR reactions amplifying specific mini– or maxicircle regions. PCR products were analyzed on agarose gels. **(D)** Growth curve of uninduced (-Tet) and induced (+Tet) bloodstream form (BSF) New York single marker (NYsm) RNAi cell line ablating MaRF11/435. Inset: RT-PCR product of MaRF11/435 mRNA in uninduced (-) or two days induced (+) cells. Tubulin mRNA serves as loading control.

### TbHslV controls the level of MaRF11

One way of how TbPam18 and TbPam16 could regulate the activity of MaRF11 would be by controlling its degradation, reminiscent to TbPIF2, the levels of which are controlled by the mitochondrial protease TbHsIVU^47^. Indeed, RNAi-mediated ablation of the TbHslV subunit of TbHslVU results in a growth retardation (*Fig. 9A*) and a concomitant accumulation of giant kDNAs as observed previously (*Fig. 9B*) ^47^. Under the same conditions, analogous to TbPIF2, a continuous increase in the levels of MaRF11 for up to six days after induction is observed, whereas the levels of TbPam16 and EF1a remained constant (*Fig. 9C*). Moreover, expression of HA-tagged MaRF11 in the induced TbPam16 RNAi cell line causes a parallel reduction on the levels of TbPam16 and MaRF11, respectively (*Fig. 9D*). This indicates that the stability of MaRF11 depends on TbPam16 and therefore likely on the presence of the TbPam16/TbPam18 heterodimer.

**Figure 9.**
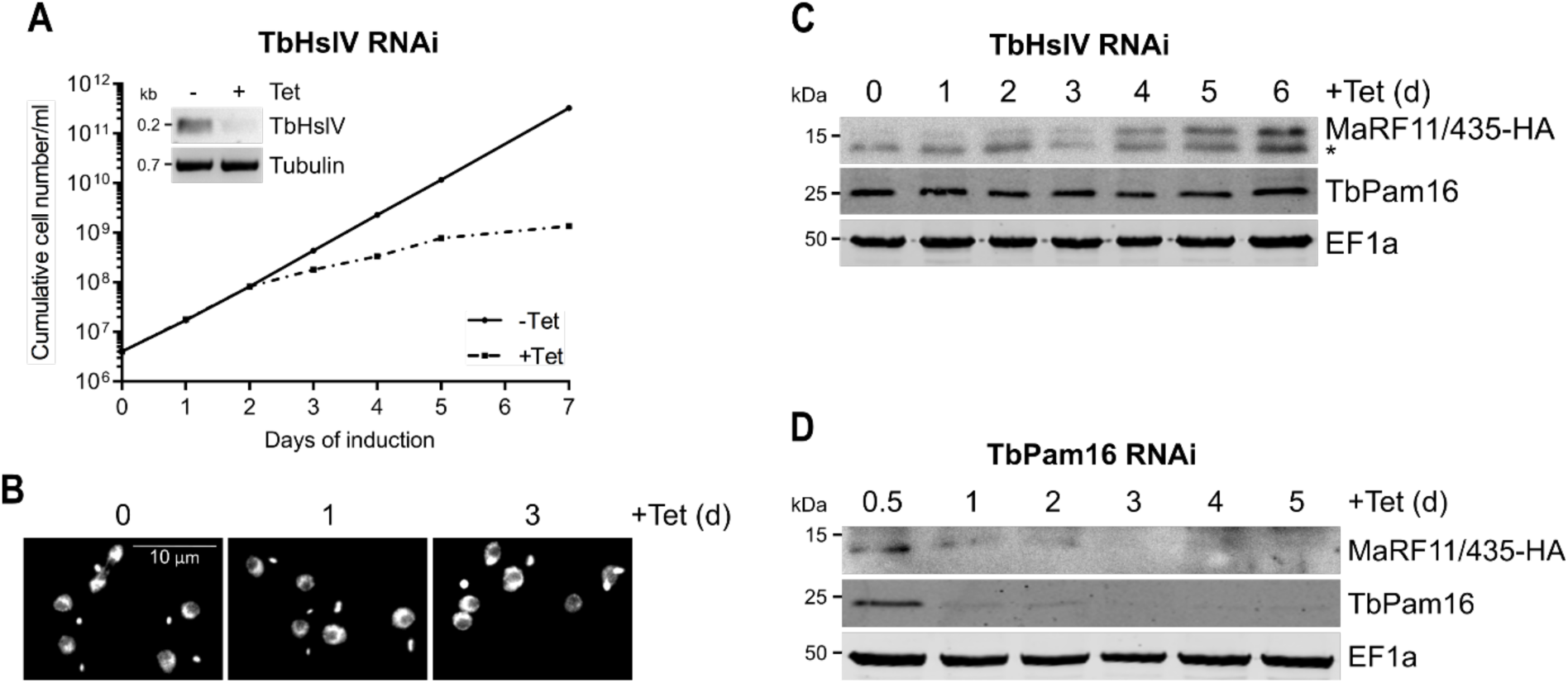
– TbHslV controls the level of MaRF11/Tb927.11.435: (A) Growth curve of uninduced (-Tet) and induced (+Tet) cells expressing MaRF11/435-HA in the background of TbHslV RNAi. Inset shows the RT-PCR product of the TbHslV mRNA in uninduced (-) or two days induced (+) cells. Tubulin mRNA serves as loading control. **(B)** Fluorescence microscopy analysis of the DAPI-stained uninduced and one or three days induced cell line explained in (A). **(C)** Immunoblot analysis of steady-state protein levels of MaRF11/435-HA and TbPam16 in TbHslV RNAi background over six days of induction. EF1a serves as loading control. Asterisk indicates an unspecific band. **(D)** Immunoblot analysis of steady-state protein levels of MaRF11/435-HA and TbPam16 in TbPam16 RNAi background between half a day and five days of induction. EF1a serves as loading control.

## Discussion

Our study has shown that TbPam18 and TbPam16 are essential for replication of the maxicircle component of the kDNA *(Fig. 2)*. This observation closes an important gap in the previously proposed evolutionary scenario explaining why trypanosomes have a single bifunctional TIM complex only^26^. Our finding is unexpected, because neither Pam18 nor Pam16 orthologues have ever been associated with the biogenesis of the mitochondrial genome before. Thus, while phylogenomics classifies TbPam18 and TbPam16 as *bona fide* Pam18 and Pam16 orthologues^26^, the two proteins have switched their function during evolution, from protein import to mitochondrial genome replication. It is worth noting that this change in function could only be discovered experimentally and was not predicted *in silico*. Interestingly, yeast Pam18 has also been associated with another function than mitochondrial protein import. It can stimulate the mHsp70-dependent assembly of respiratory super-complexes, when present as a homodimer^64^. However, in contrast to TbPam18, yeast Pam18 is still an essential component of the PAM and thus has a dual function.

Among all proteins known to be involved in kDNA maintenance and replication, depletion of only four preferentially affects the maxicircles: the mitochondrial RNA polymerase (mtRNAP)^65^, the mitochondrial DNA primase 1 (TbPRI1)^51^, the mitochondrial DNA helicase (TbPIF2)^47^ and the mitochondrial heat shock proteins TbmHsp70/TbmHsp40^52^. We have now discovered three additional factors: TbPam18, TbPam16 and the TbPam16–interacting MaRF11. Each of them is essential for normal growth of PCF *T. brucei*^26^(*Fig. 7C*) and their depletion specifically affects replication of the maxicircles prior to the onset of the growth arrest (*Fig. 2B, Fig. 8B*). Interestingly, similar to what has been observed upon ablation of the maxicircle replication factors mentioned above, total minicircle levels remain constant after the loss of maxicircles upon TbPam18, TbPam16 and MaRF11 depletion (*Fig. 2B, Fig 8B*). Moreover, depletion of TbPRI1^51^ and TbmHsp70/TbmHsp40^52^, but not of TbPIF2^47^, causes a rapid shrinkage of kDNA disks. The same was observed after TbPam18, TbPam16 and MaRF11 ablation (*Fig. 2A, Fig. 8A*). A reduction in maxicircles alone, which make up only 10% of the kDNA network, cannot explain this shrinkage. In fact, while maxicircles are selectively depleted in the TbPIF2 RNAi cell line, its kDNA network remains intact^47^. It has been suggested for TbPRI1^51^ and TbmHsp70/TbmHsp40^52^ depletion that the shrinkage of the kDNA network is due to the detachment of minicircles from the kDNA disk. The free minicircles are then replicated but cannot reattach to the maxicircle-depleted kDNA network. As a consequence, the kDNA network shrinks and free minicircles accumulate in the mitochondrial matrix^51,52^ which is exactly what is also observed in the TbPam18, TbPam16 and the MaRF11 RNAi cell lines (*Fig. 3, Fig. 8C*).

How can we explain that TbPam16, TbPam18 and MaRF11 are dispensable in the BSF of *T. brucei*? The two life cycle stages show many differences, including optimal growth temperatures (27°C for PCF/37°C for BSF) and generation times (10-12 hours for PCF/5-6 hours for BSF). Moreover, the PCF can undergo cytokinesis without completion of mitosis, whereas in the BSF a mitotic block inhibits cytokinesis but not kDNA replication^66^. Thus, it would not be surprising if these differences may require some life cycle stage specific adaptations in the regulation of kDNA replication.

TbPam18 and TbPam16 are integral IM proteins, each containing a single TMD^26^ that is essential for their function (*Fig. 5*). This contrasts with yeast, where the TMD of Pam18 is dispensable and where Pam16 does not even have a TMD^67^. Our results functionally connect maxicircle replication to the IM. This is unexpected as maxicircles are replicated while remaining interlocked with the kDNA network and proteins involved in their replication generally localize to the kDNA disk^68^. The only known integral IM protein associated with kDNA inheritance is p166, a subunit of the TAC^69,70^. The TAC is essential for kDNA segregation but not for its replication^37,71^. Knockdown of TAC subunits leads to enlarged, overreplicated kDNAs in a few cells^37,71^, which is not what is observed upon TbPam16, TbPam18 and MaRF11 depletion. It is therefore unlikely that these two proteins are involved in kDNA segregation and their IM localization must be explained in a different way.

J-domain family proteins are known regulators of a plethora of biological processes by selecting client proteins for Hsp70-type chaperones^59,72,73^. Interestingly, *T. brucei* has a greatly expanded mitochondrial J-domain protein family consisting of at least 38 members^74, 1,75–78^. One reason for this could be that they may be required for functional differentiation of the single TbmHsp70.

TbPam16, just as its yeast counterpart, is a J-like protein. However, while Pam18 in yeast contains an intact J-domain with a conserved HPD motif^79^, TbPam18 is a J-like protein containing an HSD tripeptide. Expression of a chimeric protein, in which the J-like domain of endogenous TbPam18 was replaced by the J-domain of yeast Pam18, failed to restore growth in a TbPam18 RNAi cell line (*Fig. 6AC*). Surprisingly, the change of TbPam18’s HSD to an HPD does not affect the functionality of the protein (*Fig. 6AB*). Thus, the divergence that prevents the functional interchangeability of the TbPam18 J-like domain with the J-domain of yeast Pam18 must have occurred outside the tripeptide motif.

Since the HPD motif is essential for the interaction of J-domains with their Hsp70 partners^80^, the J-like domains of TbPam18 and TbPam16 likely cannot stimulate the ATPase activity of TbmHsp70, at least not directly. Thus, the change in function of TbPam18 during evolution must have allowed the inactivation of the conserved tripeptide from HPD to HSD indicating that TbPam18 and TbPam16 function independently of mHsp70. This is reminiscent to what has been suggested for *Arabidopsis thaliana*, which has 21 J-like proteins involved in various processes, most of which function independently of Hsp70^81,82^. In line with this, pulldown experiments of TbPam16 prominently recover TbPam18, but neither TbmHsp70 nor other J-domain protein family members (*Fig 7A*). Instead, the SILAC CoIPs recovered the two essential proteins MaRF11 and TbCOQ8 indicating they might be client proteins (*Fig. 7A*). This implies that the TbPam18/TbPam16 heterodimer binds to a few clients only. Ablation of TbCOQ8 inhibits growth of PCF trypanosomes, but does not affect the size of the kDNA (*Fig. 7B*, *Fig. 8A*). However, ablation of the soluble, basic protein MaRF11 phenocopies what is observed in TbPam18 and TbPam16 RNAi cell lines (*Fig. 7B, Fig 8AB*). We have also shown that MaRF11 is a substrate of the mitochondrial proteasome orthologue TbHslVU, since depletion of the TbHslVU subunit TbHslV increases MaRF11 levels (*Fig. 9C*). Thus, both MaRF11 and the previously characterized TbPIF2, the only two TbHsIVU substrates known to date, are essential maxicircle replication factors explaining the three to four-fold accumulation of maxicircles in the absence of TbHslV^28^. Whereas the TbHsIVU substrate(s) causing the previously reported 20-fold accumulation of minicircles in the absence of TbHslVU remain unknown^28^. However, the identification of MaRF11 as a second substrate of TbHslVU underscores the central role the mitochondrial proteasome plays in controlling kDNA replication.

In *T. brucei*, the mitochondrial S-phase is precisely coordinated with the nuclear S-phase^83^. However, what signal regulates kDNA replication is not known. Based on the results describe above, we propose the following model for a membrane-bound regulatory circuit that controls the replication of the maxicircle part of the kDNA *(Fig. 10)*. This circuit comprises of four known factors: i) MaRF11, which we posit directly mediates replication, possibly by binding to maxicircles via its positive charges; ii) TbHslVU which regulates the level of MaRF11 by proteolytic degradation; and iii) The membrane-bound TbPam18/TbPam16 dimer, which binds to MaRF11 and protects it from degradation. While the presented model is compatible with our experimental findings *(Fig. 10)*, open questions remain. We do not know what role MaRF11 plays in maxicircle replication, how the binding of MaRF11 to the TbPam18/TbPam16 dimer is regulated and what the postulated nuclear signal to the proposed regulatory circuit might be. Thus, our model should serve as a guide for future experiments aimed at providing more insights into the intricate process of how the maxicircles and the kDNA in general are replicated.

**Figure 10.**
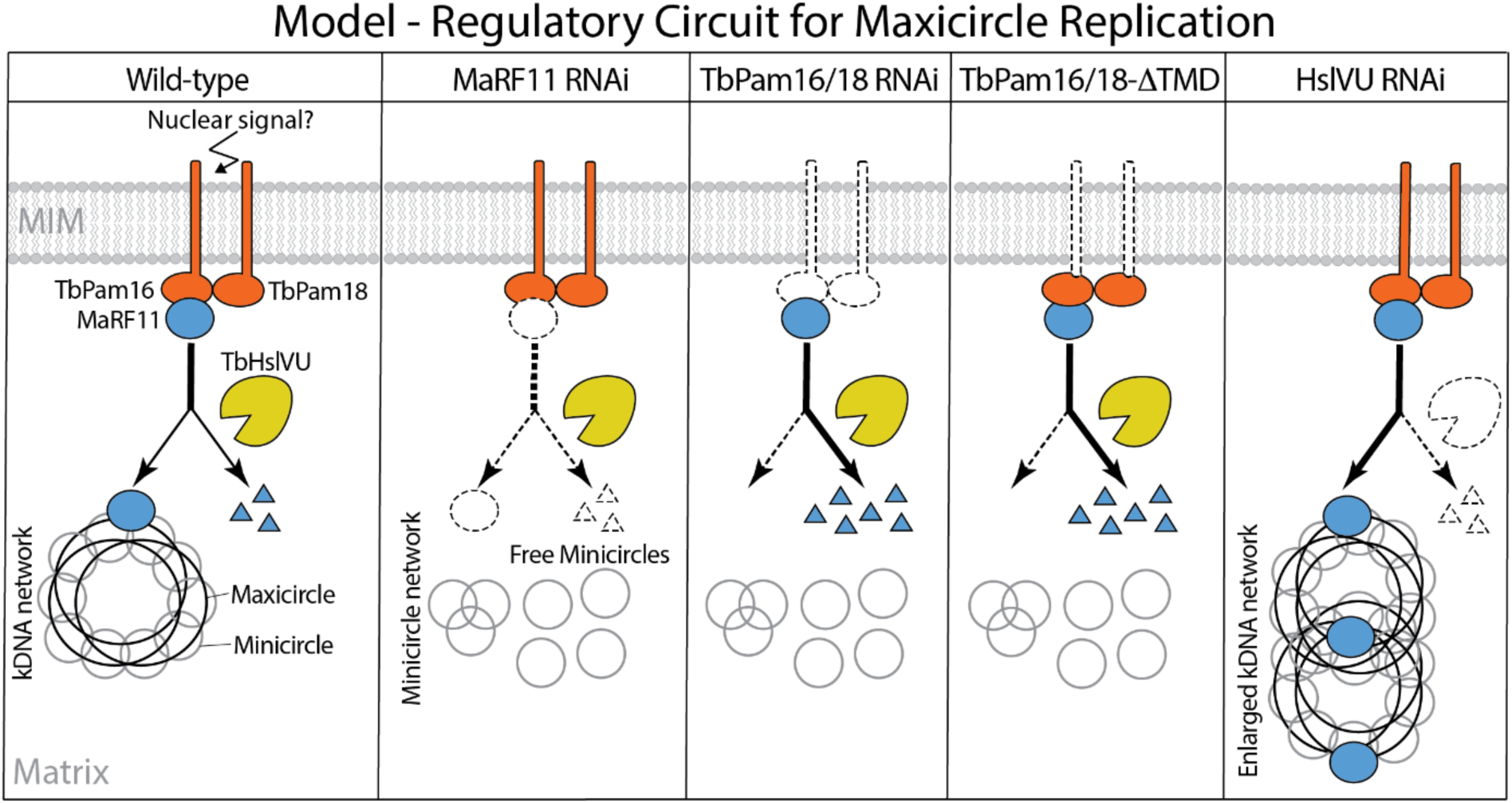
– Model of regulatory circuit for maxicircle replica_on. Wild-type: model depicting the membrane-bound regulatory circuit controlling maxicircles possibly in response to an as yet unknown nuclear signal. The circuit consists of at least four factors: TbPam18, TbPam16, MaRF11 and the mitochondrial proteasome HslVU. MaRF11 is proposed to directly mediate maxicircle replication. TbPam18 and TbPam16 protect MaRF11 from complete degradation by HslVU. MaRF11 RNAi: absence of MaRF11 depletes maxicircles and causes a shrinkage of the remaining kDNA network. Free minicircles accumulate. TbPam16/18 RNAi: depletion of either of the two proteins results in a decrease of MaRF11 levels and therefore causes the exact same phenotype as is observed aÉer MaRF11 depletion. TbPam16/18-ΔTMD: expression of TbPam16/18 lacking their transmembrane domains in the background of TbPam16/18 RNAi does not restore maxicircle copy number. HslVU RNAi: depletion of HslVU activity results in accumulation of MaRF11 and enlarged kDNA networks as observed before.Supplementary figures

## Material and methods

### Transgenic cell lines

Transgenic *T. brucei* cell lines are either based on the PCF strain 29-13 or the BSF strain NYsm^57^. PCF cells were grown in SDM-79^84^ supplemented with 10% (v/v) foetal calf serum (FCS) at 27°C. BSF cells were cultivated in HMI-9^85^ containing 10% (v/v) FCS at 37°C.

RNAi against TbPam18 (Tb927.8.6310) and TbPam16 (Tb927.9.13530) has been described previously^26^. For complementation experiments, synthetic genes (Biomatik) were used. The codons in regions of the open reading frame (ORF) that are targeted by RNAi were changed such that their transcripts are RNAi resistant (RNAi-res.) but still translate into the same amino acid sequence as in the endogenous proteins. To produce constructs allowing expression of N-terminally truncated TbPam18 (ΔN-TbPam18) and TbPam16 (ΔN-TbPam16), the corresponding DNA fragments were amplified from the synthetic genes. To ensure targeting to mitochondria, the mitochondrial targeting sequence (MTS) of trypanosomal mitochondrial Hsp60 (TbmHsp60, Tb927.10.6510) was cloned in front of the truncated constructs. For the RNAi-res. Tb/ScPam18 fusion protein and RNAi-res. TbPam18-S98P, additional synthetic genes (Biomatik) were used. To generate RNAi-res. Tb/ScPam18, the first 138 nucleotides of RNAi-res. TbPam18 were fused to the last 213 nucleotides of wildtype yeast Pam18 (YLR008C). To generate TbPam18-S98P, the cytosine at position 292 of the nucleotide sequence of RNAi-res. TbPam18 was exchanged against a thymine. Sequences of all synthetic genes are shown in *Fig. S5*.

To generate plasmids for ectopic expression of untagged, N– or C-terminally triple c– myc– or HA-tagged, RNAi-res., full-length or N-terminally truncated TbPam18 or TbPam16 variants, Tb/ScPam18, TbPam18-S98P and MaRF11 the complete or truncated ORFs of the respective genes were amplified by PCR. The PCR products subsequently were cloned into a modified pLew100 vector^57,86^, which contains a puromycin resistance cassette and either no epitope tag or triple c-myc– or HA-tags^87^.

MaRF11, TbCOQ8, TbCOQ9, Tb927.8.4190 and TbHslV (Tb927.11.10240) RNAi cell lines were generated using the same pLew100-derived vector described above. This vector allows the generation of a stem-loop construct by the insertion of the RNAi target regions in opposite directions and a 460 nucleotide (nt) spacer fragment forming the loop. The RNAi targets the indicated nts of the ORFs of MaRF11 (Tb927.11.435, nt 14-255) and TbCOQ8 (nt 353-536) or the 3’ untranslated regions (UTRs) of TbCOQ9 (nt (+49) – (+543)), Tb927.8.4190 (nt (+253) – (+458)) and TbHslV (nt (+114) – (+609)). To ensure efficient transcription of the RNAi construct in the NYsm BSF strain, the procyclin promotor in the above described MaRF11 RNAi plasmid was exchanged against the rRNA promotor.

The NYsm BSF cell line containing the TbPam16 double knockout (dKO) was generated by fusing the 500 nts up– and downstream of the TbPam16 alleles to the N– or C-terminus of the hygromycin (hygro) or blasticidin (blast) resistance cassette, respectively. The first TbPam16 allele was replaced by hygro resulting in the single KO (sKO). To generate the dKO, the second TbPam16 allele was replaced by blast.

### Antibodies

Polyclonal rabbit antiserum against TbPam16 was commercially produced (Eurogentec, Belgium) using amino acids 153-167 (VKDSHGNSRGNDAMW) as antigen. For western blots (WB), the TbPam16 antiserum was used at a 1:500 dilution. Commercially available antibodies used in this study were: Mouse anti-c-myc (Invitrogen, dilution WB 1:2’000), mouse anti-HA (Sigma-Aldrich, dilution WB 1:5’000) and mouse anti-EF1a (Merck Millipore, dilution WB 1:10’000). Polyclonal rabbit anti-ATOM40 (dilution WB 1:10’000) and polyclonal rabbit anti-Cyt C (dilution WB 1:100) were previously produced in our laboratory^78,88^.

Secondary antibodies used: Goat anti-mouse IRDye 680LT conjugated (LI-COR Biosciences, dilution WB 1:20’000) and goat anti-Rabbit IRDye 800CW conjugated (LI-COR Biosciences, dilution WB 1:20’000). For detection of 435-HA on immunoblots, HRP-coupled anti-mouse secondary antibodies (Sigma) were used.

### Digitonin extraction

Cell lines were induced with tetracycline for one day prior to the experiment to ensure expression of epitope-tagged proteins. To selectively solubilize the plasma membrane, 1 x 10^8^ cells were incubated at 4°C for 10 min in a buffer containing 0.6 M sorbitol, 10 mM Tris–HCl (pH 7.5), 1 mM EDTA (pH 8.0) and 0.015% (w/v) digitonin. A mitochondria-enriched pellet was separated from a supernatant that is enriched in cytosolic proteins by centrifugation (6’800 g, 5 min, 4°C). Equivalents of 2 x 10^6^ cells of each fraction were analysed by SDS-PAGE and western blotting.

### Alkaline carbonate extraction

A digitonin-extracted, mitochondria-enriched pellet was resuspended in 100 mM Na2CO (pH 11.5) and incubated at 4°C for 10 min. Centrifugation (100’000 g, 10 min, 4°C) yielded in a pellet enriched in integral membrane proteins and a supernatant enriched in soluble or loosely membrane-associated proteins. Equivalents of 2 x 10^6^ cells of each fraction were analysed by SDS-PAGE and western blotting.

### Fluorescence microscopy and kDNA area quantification

TbPam18 and TbPam16 RNAi cells were fixed with 4% paraformaldehyde in PBS, postfixed in cold methanol and mounted using VectaShield containing 4ʹ,6-diamidino-2-phenylindole (DAPI) (Vector Laboratories). Images were acquired by a DMI6000B microscope and a DFC360 FX monochrome camera (both Leica Microsystems).

Images were analysed using ImageJ^89^. The kDNA size analysis was performed on binarized 8-bit format images. The size of particles was measured in arbitrary units (a.u.) and kDNA particles >0.0 a.u. and <0.75 a.u. were included in the analysis. Boomerang shaped, dividing kDNAs and randomly picked up particles were manually removed from the analysis. Significance of these results was calculated by an unpaired two-tailed t-test.

### RNA extraction, RT-PCR and northern blotting

Acid guanidinium thiocyanate-phenol-chloroform extraction to isolate total RNA from uninduced and two days induced RNAi cells was done as described elsewhere^90^. To determine RNAi efficiency, the extracted RNA was either utilized for RT-PCR or separated on a 1% agarose gel in MOPS buffer containing 0.5% formaldehyde for subsequent northern blotting. Northern probes were generated from gel-purified PCR products corresponding to the RNAi inserts or the overexpressed proteins described above, and radiolabelled by means of the Prime-a-Gene labelling system (Promega).

### DNA extraction, Southern blotting and quantitative PCR

For DNA isolation, 5 x 10^7^ cells were resuspended in NTE buffer (100 mM NaCl, 10 mM Tris (pH 7.5) and 5 mM EDTA) containing 0.5% SDS for cell lysis and 0.2 mg/ml RNase A to degrade RNA. After incubation for 1 hr at 37°C, 1 mg/ml proteinase K was added, followed by 2 hr of incubation at 37°C. DNA was isolated by phenol-chloroform extraction and subsequent ethanol precipitation.

For Southern blotting, 5 μg of DNA were digested overnight at 37°C with HindIII and XbaI. Digested DNA was separated in a 1% agarose gel in 1X TAE buffer. Gel processing and blotting was done as described elsewhere^50,91^. For kDNA detection, sequence-specific mini– and maxicircle probes were generated by PCR. The minicircle probe was a 0.1 kb stretch of the conserved minicircle sequence^91^. A 1.4 kb fragment served as the maxicircle probe^50,92^. For normalization, a tubulin probe binding to a 3.6 kb stretch within the intergenic region between α– and β-tubulin, was used^91^. Probes were radiolabelled by means of the Prime-a-Gene labelling system (Promega).

To determine total mini– and maxicircles or free minicircle levels by quantitative PCR, DNA was either isolated from whole cells or from fractionated digitonin-extracted, mitochondria-enriched pellets and used as the template in a PCR utilizing the same primers as for the Southern blot probes.

### SILAC RNAi and SILAC CoIP experiments

TbPam18 and TbPam16 RNAi cells or cells exclusively expressing TbPam16-HA were washed in PBS and resuspended in SDM-80^93^ containing 5.55 mM glucose, 10% dialyzed FCS (BioConcept, Switzerland) and either light (^12^C6/^14^Nχ) or heavy (^13^C6/^15^Nχ) isotopes of arginine (1.1 mM) and lysine (0.4 mM) (Euroisotope). The cells were grown in SILAC medium for six to ten doubling times to ensure a complete labelling of all proteins with heavy amino acids. For the SILAC RNAi and the premix SILAC CoIP, uninduced and induced (four days for SILAC RNAi, two days for SILAC CoIP) cells were mixed in a one-to-one ratio. For the postmix SILAC CoIP, uninduced and induced cells were kept separately. From all samples, digitonin-extracted, mitochondria-enriched pellets were generated. For the SILAC RNAi experiments, these pellets were processed as described previously including tryptic in solution digestion^94^ and then analysed by liquid chromatography-mass spectrometry (LC-MS). TbPam18 and TbPam16 SILAC RNAi experiments were done in four biological replicates including a label-switch.

For the SILAC CoIP experiments, mitochondria-enriched digitonin pellets were solubilized in a buffer containing 20 mM Tris-HCl (pH 7.4), 0.1 mM EDTA, 100 mM NaCl, 10% glycerol, 1X Protease Inhibitor mix (Roche, EDTA-free) and 1% (w/v) digitonin for 15 min at 4°C. After centrifugation (21’000 g, 15 min, 4°C), the lysate was transferred to HA bead slurry (anti-HA affinity matrix, Roche), which had been equilibrated in wash buffer (20 mM Tris-HCl (pH 7.4), 0.1 mM EDTA, 1 mM NaCl, 10% glycerol, 0.2% (w/v) digitonin). Incubation in an end-over-end shaker for 2 hr at 4°C was followed by removal of the supernatant containing the unbound proteins. After washing the bead slurry three times with wash buffer, the bound proteins were eluted by boiling the resin for 5 min in 2% SDS in 60 mM Tris-HCl (pH 6.8). In case of the postmix SILAC CoIP, eluates of uninduced and induced cells were now mixed in a one-to-one ratio. All eluates were further prepared for analysis by LC-MS as has been described in detail elsewhere^95^. TbPam16-HA pre– and postmix SILAC CoIP experiments were done in four biological replicates including label-switches.

### LC-MS and data analysis

LC-MS analyses of tryptic peptide mixtures from all experiments were performed using a Q Exactive Plus mass spectrometer connected to an UltiMate 3000 RSLCnano HPLC system (Thermo Fisher Scientific, Germany) as described before^26^ with minor modifications. The software package MaxQuant ^96,97^ (versions 1.6.3.4 and 2.0.2.0 for SILAC RNAi and SILAC CoIP data, respectively) was used for protein identification and SILAC-based relative quantification. Mass spectrometric raw data were searched against a database containing the protein sequences for *T. brucei* TREU927 as provided by the TriTrypDB (https://tritrypdb.org; version 8.1). Protein identification and quantification was based on ≥ 1 unique peptide and ≥ 1 ratio count, respectively. For all other parameters, MaxQuant default settings were used, including carbamidomethylation of cysteine as fixed modification, N-terminal acetylation and oxidation of methionine as variable modifications, and Lys8/Arg10 as heavy labels. The options “requantify” and “match between runs” were enabled.

MaxQuant result files were processed with python using pandas (version 1.5.3; https://pandas.pydata.org) as well as numpy (version 1.24.2; https://numpy.org/), seaborn (version 0.11.2; https://seaborn.pydata.org), scipy (version 1.10.0; https://www.scipy.org/), and matplotlib (version 3.6.3; https://matplotlib.org/) for data analysis and visualization.

Data analysis of all experiments was based on protein abundance ratios calculated by MaxQuant.

To identify proteins affected by ablation of TbPam16 and TbPam18 following RNAi induction, MaxQuant protein ratios were first normalized replicate-wise by adjusting the summed ratios to the highest value, followed by cyclic loess normalization^98^ of log2-transformed protein ratios as implemented in the R Bioconductor (version 3.17) package “affy” ^99^ (version 1.78.2). Values missing in one or two out of four replicates were imputed using the DIMA package ^100^ (https://github.com/kreutz-lab/DIMAR). To identify proteins with significantly altered abundance upon RNAi induction, the “linear models for microarray data” (limma) approach)^101,102^ (version 3.28.14) was applied. *P* values were corrected for multiple testing following the Benjamini-Hochberg method^103^.

To identify proteins significantly enriched in TbPam16 pre– and postmix SILAC CoIP experiments, the rank sum method^104,105^ as implemented in the R package “RankProd” ^106^ (version 3.24.0) was applied. The rank sum, defined as the arithmetic mean of the ranks of a protein in all replicates, was converted into a *P* value and a false discovery rate. For information about proteins identified and quantified, see *Supplementary Table S1* (SILAC RNAi experiments) and *Supplementary Table S2* (SILAC IP experiments).

## Data availability

The mass spectrometry proteomics data have been deposited to the ProteomeXchange Consortium^107^ via the PRIDE^108^ artner repository and are accessible using the dataset identifiers PXD046840 (TbPam16 SILAC RNAi data), PXD046845 (TbPam18 SILAC RNAi data), and PXD046849 (TbPam16 SILAC CoIP data).

## Acknowledgements

Work in the lab of A.S. was supported in part by NCCR RNA & Disease, a National Centre of Competence in Research (grant number 205601) and by project grant SNF 205200 both funded by the Swiss National Science Foundation. We thank Julian Bender and Johannes Zimmermann for assistance in bioinformatics data analysis.

## Supplementary figures

**Figure S1.**
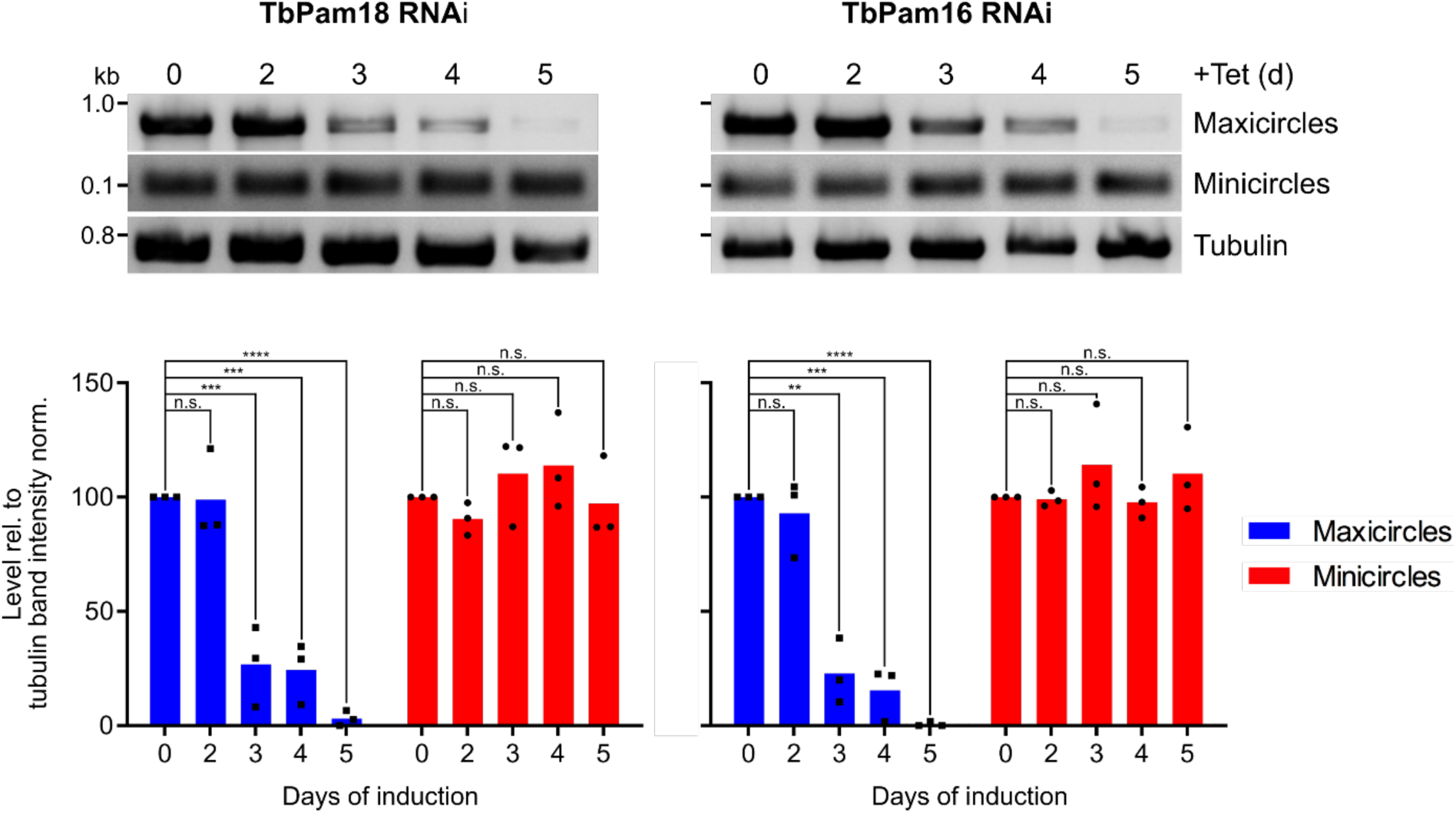
– Changes in mini– and maxicircle levels can be detected by quan_ta_ve PCR: Upper panels: A quantitative PCR-based (qPCR) method was used to detect changes in steady-state levels of total mini– and maxicircles. Total DNA was extracted from uninduced and two to five days induced TbPam18 and TbPam16 RNAi cell lines. This DNA was used as the template in PCRs amplifying specific mini– and maxicircle regions or the intergenic region of tubulin. PCR products were analyzed on agarose gels. Lower panels: Densitometric quantification of mini– and maxicircle abundance as detected by qPCR. The ratio of the mini– or maxicircle band intensity and the respective control (tubulin band intensity) was normalized (norm.) to the ratios of uninduced cells. Blue (maxicircles) and red (minicircles) bars represent the mean of three independent biological replicates. n.s.: not significant, **: *P* value<0.05, ***: *P* value<0.005, ****: *P* value<0.0001, as calculated by an unpaired two-tailed t-test.

**Figure S2.**
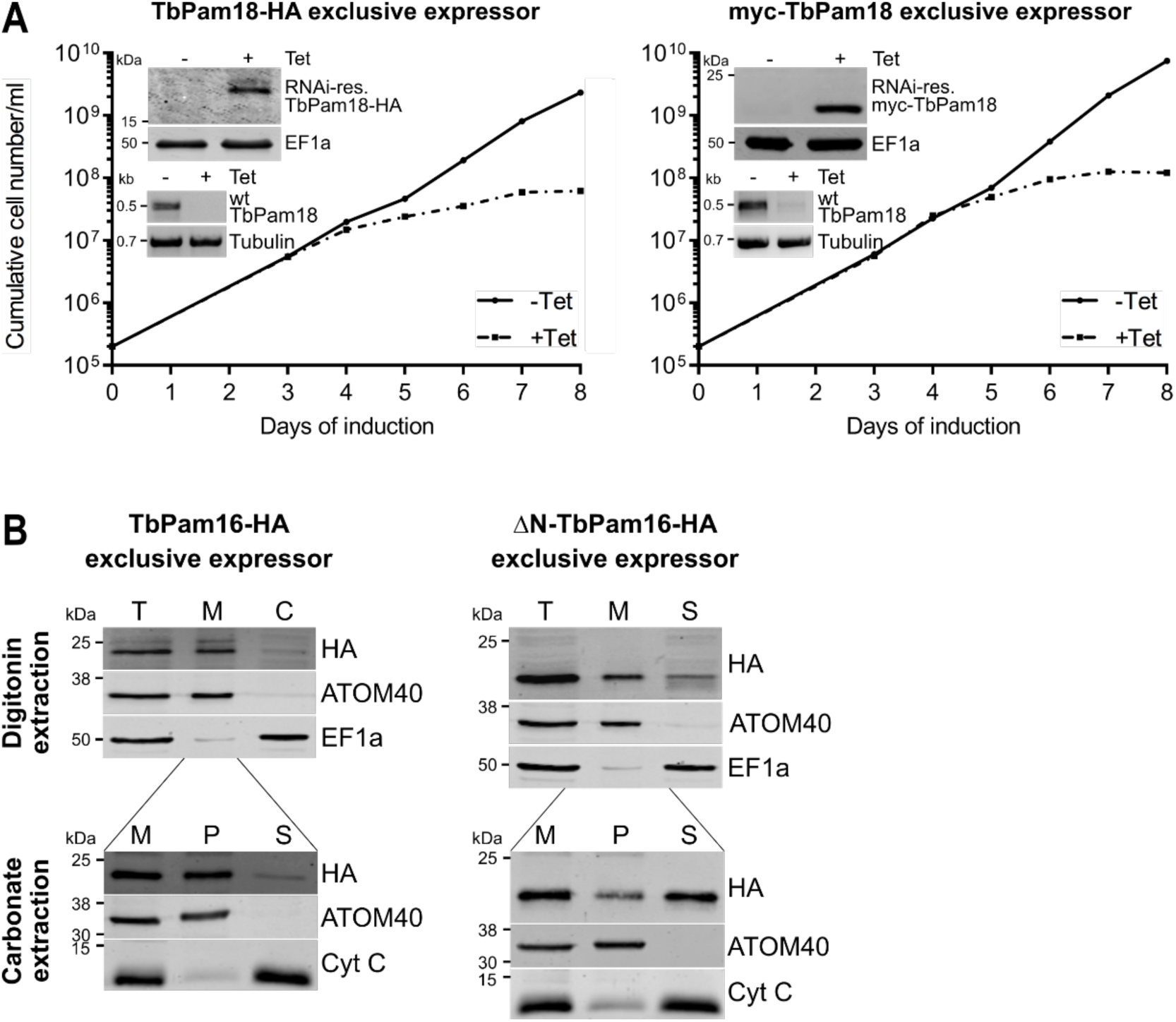
– Func_onal analysis of tagged TbPam18 and subcellular localiza_on of TbPam16 variants: (A) Growth curve of uninduced (-Tet) and induced (+Tet) cells expressing RNAi-resistant (RNAi-res.) TbPam18-HA (leÉ) or myc-TbPam18 (right) in the background of RNAi against the wildtype (wt) TbPam18 (TbPam18-HA and myc-TbPam18 exclusive expressors). Insets, top: Immunoblot analysis of whole cell-extracts of uninduced (-) and two days induced (+) cells, probed for RNAi-res. TbPam18-HA or myc-TbPam18 and EF1a as loading control. Insets, boÖom: RT-PCR products of the wt TbPam18 mRNA in uninduced (-) or two days induced (+) cells. Tubulin mRNA serves as loading control. **(B)** Upper panels: Immunoblot analysis of total cells (T), digitonin-extracted, mitochondria-enriched (M) and soluble cytosolic (S) fractions of TbPam16-HA and ΔN-TbPam16-HA exclusive expressor cell lines. Blots were probed with anÇ-HA antibodies and antisera against ATOM40 and EF1a, which serve as mitochondrial and cytosolic markers, respectively. Lower panels: Digitonin-extracted, crude mitochondrial fractions (M) were subjected to an alkaline carbonate extraction resulting in a pellet enriched in integral membrane proteins (P) and a soluble supernatant fraction (S). Immunoblots were probed with anÇ-HA and antisera against ATOM40 and cytochrome C (Cyt C), which serve as marker for integral membrane and soluble proteins, respectively.

**Figure S3.**
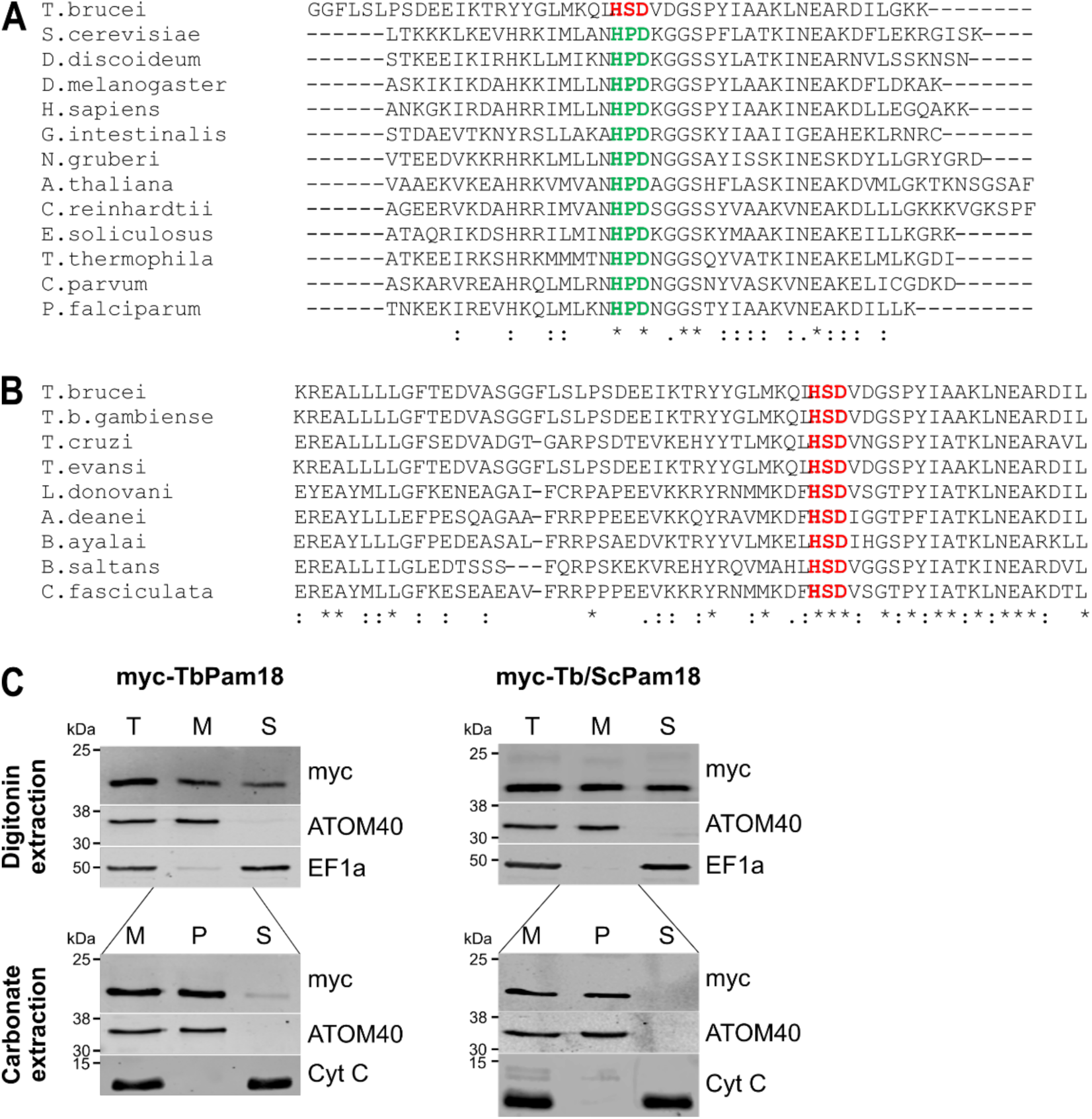
– Mul_ple sequence alignments of Pam18 homologues and subcellular localiza_on of TbPam18 variants: (A) Sequence alignment of N-terminal regions of Pam18 homologues of 13 representative eukaryotes. **(B)** Sequence alignment of N-terminal regions of Pam18 homologues of nine representative trypanosomatids. In (A) and (B) Histidine-Proline-Aspartate (HPD) motifs are highlighted in red and Histidine-Serine-Aspartate (HSD) motifs in green. **(C)** Upper panels: Immunoblot analysis of total cells (T), digitonin-extracted mitochondria-enriched (M), and soluble cytosolic (S) fractions of cell lines expressing N-terminally myc-tagged TbPam18 or Tb/ScTbPam18. Blots were probed with anÇ-myc antibodies and antisera against ATOM40 and EF1a, which serve as mitochondrial and cytosolic markers, respectively. Lower panels: Digitonin-extracted crude mitochondrial fractions (M) were subjected to an alkaline carbonate extraction resulting in a pellet enriched in integral membrane proteins (P) and a soluble supernatant fraction (S). Immunoblots were probed with anÇ-myc and antisera against ATOM40 and Cyt C, which serve as makers for integral membrane and soluble proteins, respectively.

**Figure S4.**
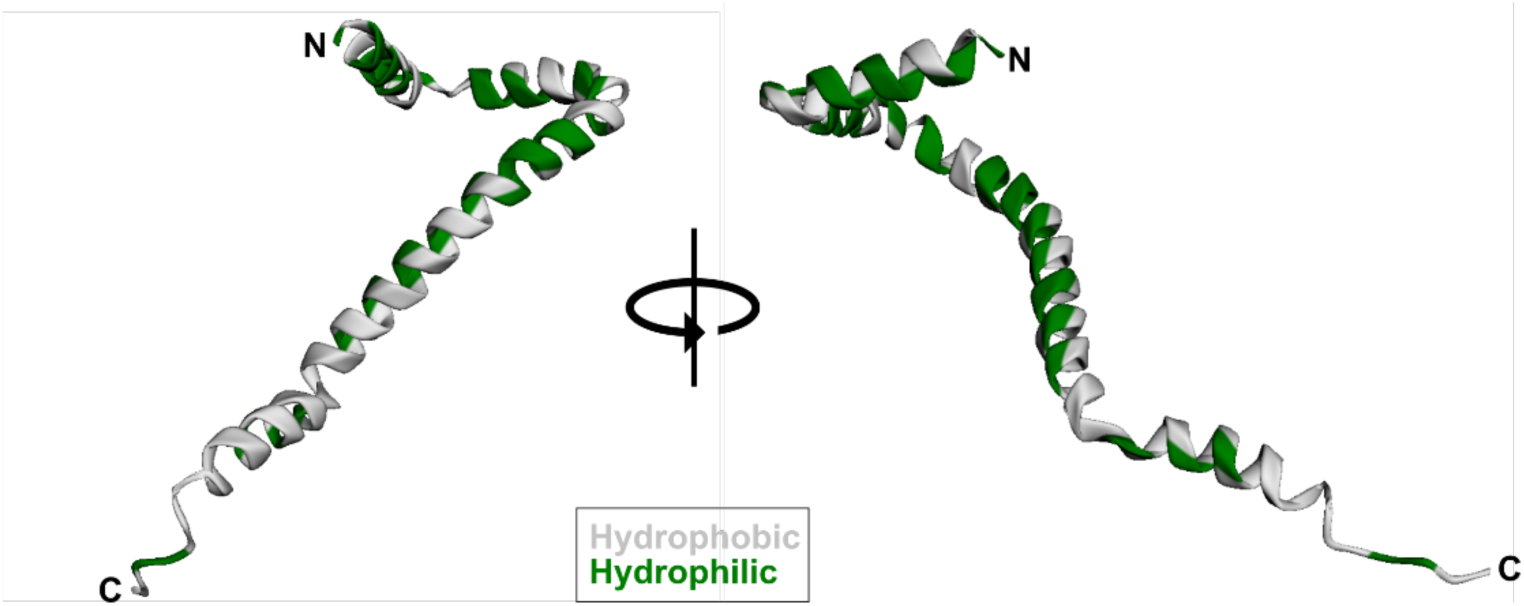
– AlphaFold predic_on of MaRF11/Tb927.11.435.

**Figure S5.**
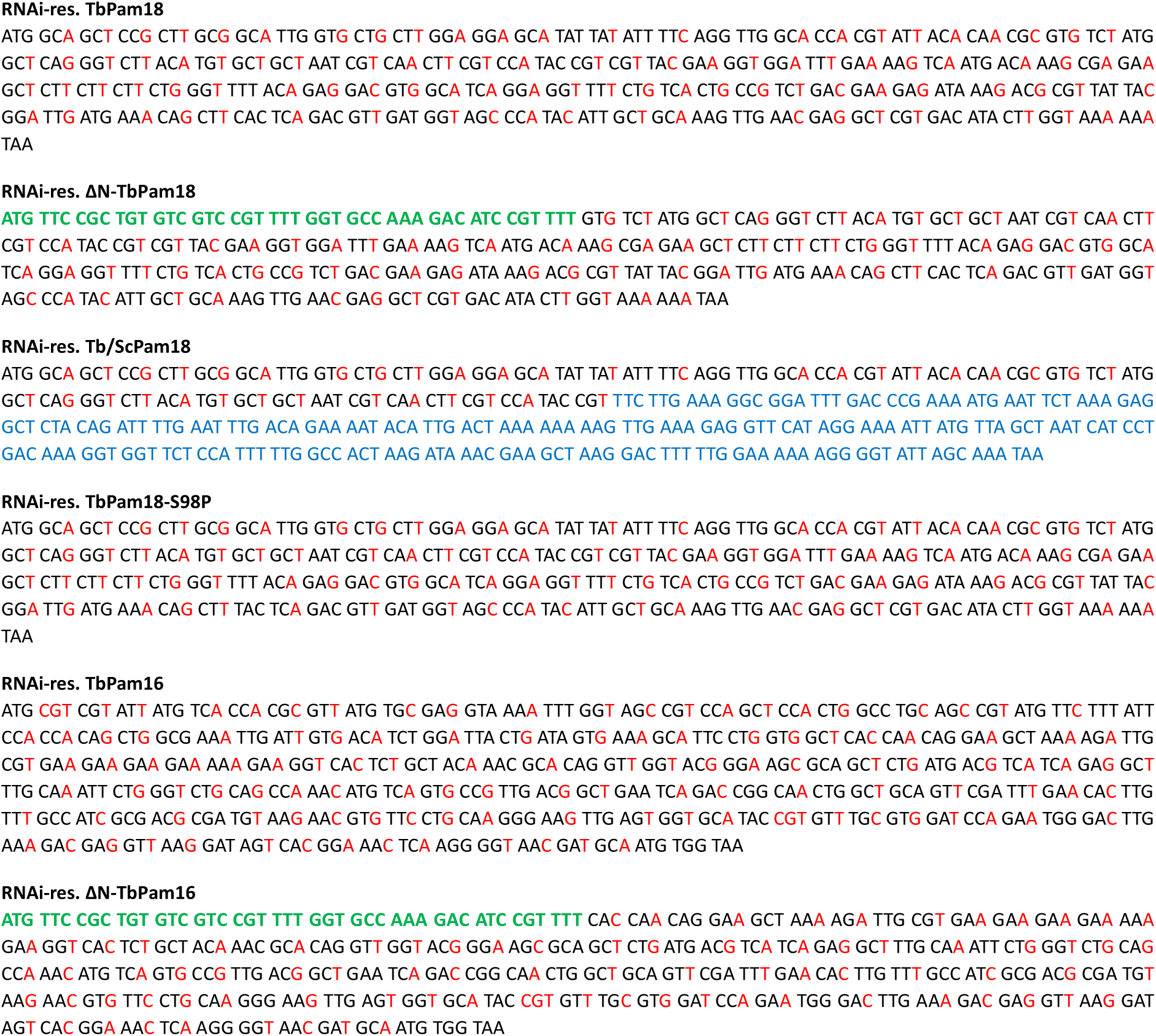
– Synthe_c TbPam18 and TbPam16 genes: RNAi-resistant (RNAi-res.) TbPam18 and TbPam16 DNA sequences in black with changed nucleotides highlighted in red. In the ΔN-TbPam18 and ΔN-TbPam16 constructs, the first 81 and 156 nucleotides, respectively, were replaced by the first 45 nucleotides of TbmHsp60, which encode the mitochondrial targeting sequence of the protein (green). To generate the Tb/ScPam18 fusion protein, the first 295 nucleotides of RNAi-res. TbPam18 were fused to the last 213 nucleotides of yeast (Sc) Pam18 (blue). To generate RNAi-res. TbPam18-S98P, the cytosine at position 292 of the nucleotide sequence was exchanged against a thymine (highlighted in yellow).

